# Dynamic subtype- and context-specific subcellular RNA regulation in growth cones of developing neurons of the cerebral cortex

**DOI:** 10.1101/2023.09.24.559186

**Authors:** Priya Veeraraghavan, Anne K. Engmann, John J. Hatch, Yasuhiro Itoh, Duane Nguyen, Thomas Addison, Jeffrey D. Macklis

## Abstract

Molecular mechanisms that cells employ to compartmentalize function via localization of function-specific RNA and translation are only partially elucidated. We investigate long-range projection neurons of the cerebral cortex as highly polarized exemplars to elucidate dynamic regulation of RNA localization, stability, and translation within growth cones (GCs), leading tips of growing axons. Comparison of GC-localized transcriptomes between two distinct subtypes of projection neurons– interhemispheric-callosal and corticothalamic– across developmental stages identifies both distinct and shared subcellular machinery, and intriguingly highlights enrichment of genes associated with neurodevelopmental and neuropsychiatric disorders. Developmental context-specific components of GC-localized transcriptomes identify known and novel potential regulators of distinct phases of circuit formation: long-distance growth, target area innervation, and synapse formation. Further, we investigate mechanisms by which transcripts are enriched and dynamically regulated in GCs, and identify GC-enriched motifs in 3 ’ untranslated regions. As one example, we identify *cytoplasmic adenylation element binding protein 4* (CPEB4), an RNA binding protein regulating localization and translation of mRNAs encoding molecular machinery important for axonal branching and complexity. We also identify *RNA binding motif single stranded interacting protein 1* (RBMS1) as a dynamically expressed regulator of RNA stabilization that enables successful callosal circuit formation. Subtly aberrant associative and integrative cortical circuitry can profoundly affect cortical function, often causing neurodevelopmental and neuropsychiatric disorders. Elucidation of context-specific subcellular RNA regulation for GC- and soma-localized molecular controls over precise circuit development, maintenance, and function offers generalizable insights for other polarized cells, and might contribute substantially to understanding neurodevelopmental and behavioral-cognitive disorders and toward targeted therapeutics.

## Introduction

Neurons are extreme examples of polarized cells, with highly specialized subcellular compartments^3-8^. The spatial enrichment of specific transcripts to distal compartments, such as axons and dendrites, is likely linked to targeted RNA trafficking based on a combination of specific motifs^9-15^ and differential expression of RNA binding proteins (RBPs)^16-20^, reviewed in^22^, as well as adjustments of RNA stability and local turnover mediated by RBPs and miRNAs^23,24^, reviewed in^25^. During development, axonal GCs navigate complex extracellular environments by responding to diverse substrate-bound and diffusible guidance cues. Each subtype-specific GC sequentially alters and refines its trajectory and growth rate, under control of such attractant and repellent cues, bypassing and rejecting inappropriate targets, until it reaches its correct distant target area, where it ultimately matures into a subtype-specific synapse (reviewed in^26^). Axonal GCs typically function in guidance 10^2^ - 10^4^ cell body diameters away from their parent somata, then function in target selection and synapse formation/maintenance 10^3^ – 10^5^ cell body diameters away. Thus, signaling to and from the nucleus via fast axonal transport would typically take hours to days^27^, while GC responses are known to occur in seconds to minutes^28^. It is known that local translation of at least some transcripts within the axon compartment is critically involved in axon guidance^29-31^, synapse formation^32,33^, maintenance^34,35^, plasticity^36,37^, and regeneration^38-43^. Further, research from our laboratory and others has identified striking spatial segregation of RNA and protein molecular machinery into distinct local subcellular domains in projection neurons (e.g., GCs and axons *vs*. parent somata). Due to technical challenges, molecular compositions of GCs have been mostly studied *in vitro* (e.g., ^6,44-46^), and *in vivo* data have been limited^5,47-49^. Our laboratory recently developed a set of experimental and analytic approaches enabling direct quantitative comparison of GC and soma RNA and protein molecular machinery in a subtype- and stage-specific manner directly from projection neurons in mouse brain^7,50^.

Immensely diverse subtypes of cortical projection neurons form connections with similarly diverse target areas throughout the central nervous system (CNS), establishing exquisitely precise functional circuitry that enables sensory integration, motor processing, associative behaviors, and advanced cognition. Callosal projection neurons (CPN) connect across the corpus callosum to the contralateral cortical hemisphere, while subcerebral (SCPN) and corticothalamic projection neurons (CThPN) extend their axons out of cortex, through the internal capsule, and beyond to innervate diverse subcerebral structures in the midbrain, brainstem, and spinal cord (for SCPN) or thalamus (for CThPN)^51-55^. Work from many labs over the past two decades provides an increasingly detailed understanding how a few best-studied subtypes of cortical projection neurons are specified by combinatorial transcriptional regulators through early steps of corticogenesis^53,55,56^. In sharp contrast, it is not well understood how distinct subtypes of projection neurons establish precise function-specific circuitry during development. Beyond axon guidance and establishment of anatomical connectivity, it is also poorly understood how subtype-specific synapses emerge from these processes to support specific functions. These questions are particularly relevant in the context of miswiring of cortical circuitry. Thus, they are closely associated with the etiology of neurodevelopmental and neuropsychiatric diseases, including autism spectrum disorders (ASD)^57,58^ and schizophrenia^59,60^.

Here, we employ our recently developed subcellular approaches to investigate whether and how spatial distribution of transcripts between GCs and their parent somata might differ between distinct subtypes of projection neurons, and how it is dynamically regulated across distinct stages of axon development. First, we identify that a large set of shared GC-localized molecular machinery is common between subtypes of cortical projection neurons, and that GC-localized transcriptomes are enriched for genes associated with neurodevelopmental and neuropsychiatric disorders. For example, we identify that transcripts of several core components and regulatory elements of the wave regulatory complex (WRC) are locally enriched in GCs of CPN and CThPN. Previous reports have linked this complex to axonal fasciculation, growth, and guidance in projection neurons^61-64^. Localization of a pool of WRC-associated transcripts to PN GCs raises the intriguing possibility of targeted local translation, assembly, and activity-dependent fine-tuning of this multimeric regulatory system for actin nucleation, assembly, and branching. Second, subtype-specific transcripts that are differentially abundant in GCs of either CPN or CThPN at postnatal day 3 (P3) are functionally enriched for classes of genes involved in either axonal growth, guidance, and synapse formation (for CPN) or translation (for CThPN). Third, intriguingly, we identify many genes with no substantial transcriptional distinction between CPN and CThPN somata, but for which GC enrichment differs sharply between subtypes at P3. Investigation of 3 ’ untranslated regions (UTRs), known to be involved with RNA trafficking and stability, identifies a set of motifs that are enriched in GC-localized transcripts of both subtypes. In particular, we identify that cytoplasmic polyadenylation elements (CPEs) are enriched in 3 ’UTRs in GCs. We further identify the RBP CPEB4 as an exemplar that likely mediates aspects of subcellular RNA localization, stability, and translation, and that dysregulates axonal branching and complexity when perturbed. Lastly, we extend this work to investigate dynamic changes of subcellular RNA localization during development within CPN, and identify motifs in 3 ’UTRs of GC-localized transcripts that are enriched in GCs during early (axon elongation and branching/collateralization) or later stages (early synapse formation) of development. These differentially enriched motifs, coupled with evidence of differential GC-localization of transcripts, strongly suggest stage-specific developmental programs for anterograde RNA trafficking, subcellular localization, and stability. We identify the RBP RBMS1 as an exemplar that regulates developmental stage-specific, dynamic changes in composition of the CPN-specific GC-localized transcriptome, required for proper CPN circuit formation.

## Results

### Cortical projection neurons contain distinct local transcriptomes in spatially segregated and functionally specialized subcellular compartments

To investigate subcellular molecular machinery that might be critical for subtype-specific circuit generation, we directly compared subcellular transcriptomes of CPN and CThPN. We isolated somata and GCs in parallel from these two distinct subtypes of cortical projection neurons directly from the early postnatal mouse brain (Figure 1a). These cortical projection neuron subtypes have their somata in distinct cortical layers, and they generate distinct circuitry (Figure 1a). CPN are central for sensory-motor integration, and for integrative and associative behavior and cognition. Their projections are predominantly intratelencephalic, extending their axons across the midline via the corpus callosum to innervate distinct bimodal homotopic target regions in the contralateral hemisphere^65-69^. CThPN are central for multi-modal sensory and motor integration and modulation between cortex and thalamus. They respect the midline, and project their axons into the internal capsule to form precise and modality-specific reciprocal circuitry with thalamocortical projection neurons in the various thalamic nuclei (reviewed in^70^).

**Figure 1:**
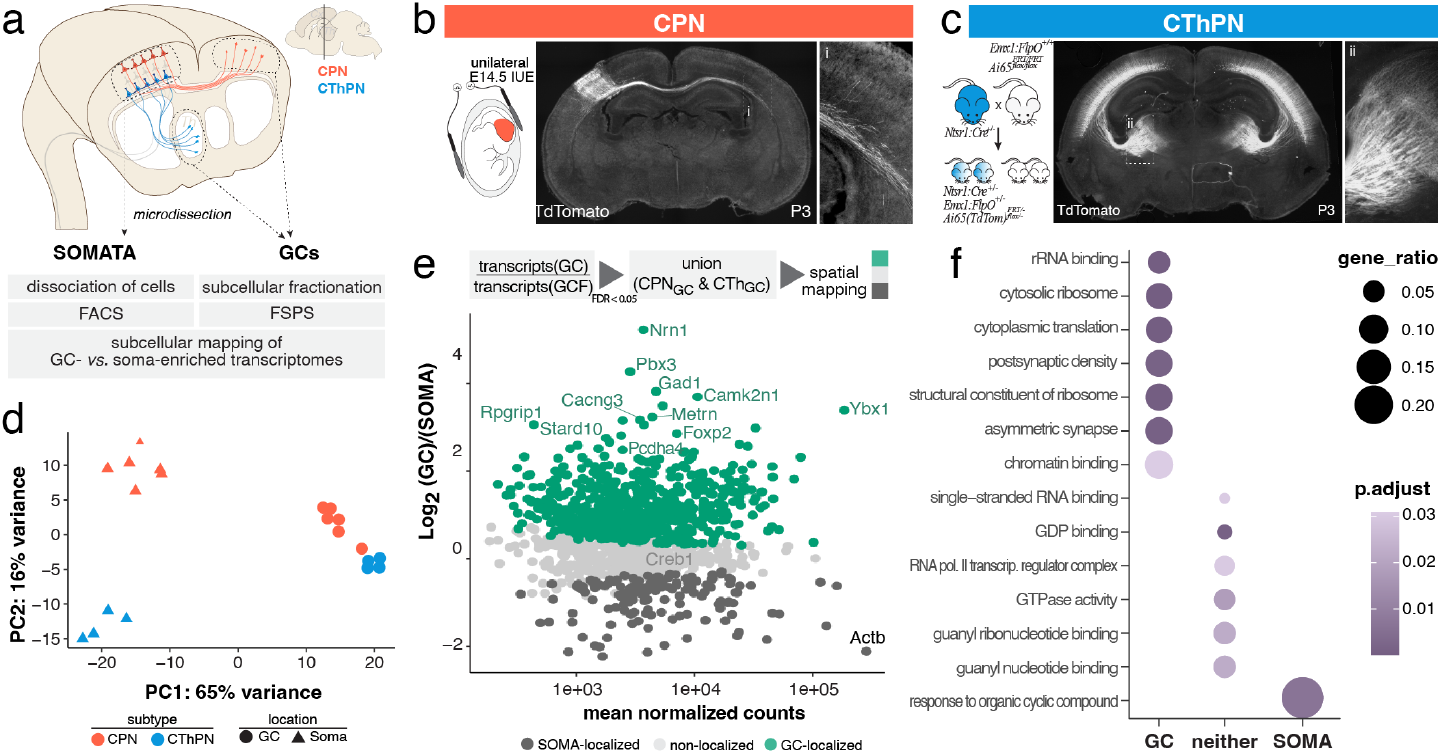
Subtypes of cortical projection neurons contain spatially segregated transcriptomes poised for subcellular, compartment-specific function. **(a)** Somata and GCs of callosal projection neurons (CPN, orange) and corticothalamic projection neurons (CThPN/CTh, blue) were isolated from mouse brains, and purified in parallel by fluorescence-based cell and small particle sorting, respectively. Subcellular RNA was isolated, and transcriptomes were quantitatively assessed, from subtype-specific GCs and somata to investigate spatial distribution of transcripts. **(b)** CPN in cortical layer II/III were labeled by E14.5 unilateral *in utero* electroporation. At P3, labeled CPN axons cross the midline and innervate homotopic targets in the contralateral cortex (inset i). **(c)** CThPN of cortical layer VI were labeled using intersectional mouse genetics (Ntsr1::Cre, Emx1::FlpO, Ai65). At P3, labeled CThPN axons have traversed the internal capsule and are innervating the thalamus (inset ii). **(d)** Principal component analysis reveals tight clustering and distinction of data points for the same subcellular compartment of either subtype (CPN; CThPN) and separation between compartments (GCs *vs*. somata) by the first component. The second component further separates samples according to subtypes. **(e)** Within the union of ambient corrected, GC-enriched transcripts across subtypes, the majority of transcripts are preferentially GC-localized (green), with smaller fractions showing no preferential subcellular localization (light grey), or preferential localization to the somata (dark grey). **(f)** Distribution of biological function assessed by GO term enrichment across GC and soma compartments. Size corresponds to the proportion of genes in the gene set (x axis) that are also in the GO term set (y axis); color corresponds to adjusted FDR.

We targeted layer II/III CPN (the dominant CPN population) for parallel GC and soma isolation using unilateral *in utero* electroporation at E14.5 with a multi-cistronic DNA construct encoding membrane-targeted (for GCs) and nuclear-localized (for somata) fluorescent reporters, resulting in reproducible labeling of SatB2-positive neurons in layers II/III of lateral somatosensory cortex (Figure 1b, Supplementary Figure 1a-c).

CThPN were targeted specifically using an intersectional transgenic approach (Emx1::FlpO, Ntsr1::Cre, Ai65), resulting in selective labeling of neurons in cortical layer VI (Figure 1c), colocalized with the widely utilized CThPN marker TBR1, and negative for the CPN marker SatB2 (Supplementary Figure 1d). Specificity of each labeling approach for the two distinct neuronal subtypes is confirmed by transcriptomic expression data of well-established marker genes (Supplementary Figure 1e). Importantly, tissue containing GCs of interest did not contain any labeled cell bodies or dendrites, which might otherwise potentially contaminate the purified GC samples (Figure 1b). Subcellular-specific GCs and somata were isolated in parallel directly from the mouse brain, using subcellular fractionation coupled with fluorescent small particle sorting (FSPS; Supplementary Figure 2a-e,^7,50^) and established fluorescence activated cell sorting (FACS) techniques^51,71^, respectively. Subcellular fractionation resulted in enrichment of GCs and distal axon particles (growth cone fraction, GCF), which contained sealed membranous spheres protecting RNA and protein content in RNase and protease hydrolysis protection assays, and showed enrichment for distal axon markers (e.g. GAP43) and de-enrichment of somatic and dendritic markers (e.g. Gm130, Map2) in western blot analysis (Supplementary Figure 2f-h). Libraries were prepared from the extracted polyA+ mRNA using SMART-Seq v4 chemistry and sequenced to a depth of >40M reads per sample (workflow outlined in Supplementary Figure 3). Most reads mapped to coding sequences within the nuclear genome (Supplementary Figure 3c/d). In GC samples, in particular, reads displayed higher bias to 3 ’ end coverage than in somata (Supplementary Figure 3e). To account for potential ambient contaminating RNAs that originate from the environment around GCs, we compared transcriptomes obtained from sorted GCs to those obtained from the GCF input and identified 865 CPN_GC_ and 624 CThPN_GC_ genes enriched in the FSPS-purified GCs over the GCF input, which we term “GC genes” (Supplementary Figure 4a & b). This bioinformatic filter removes reads that can be attributed to non-neuronal cell types or functions (Supplementary Figure 4c & d).

We first focused on transcriptome comparisons between GC and soma subcellular compartments at P3. At the level of the whole transcriptome, samples from distinct subtypes and subcellular locations clearly separate via principal component analysis, with the subcellular axis dominating the variance observed (Figure 1d). We next compared GC genes found in either subtype by their relative abundance in GCs compared to somata (Figure 1e) and identify that genes most significantly enriched in GCs are enriched for gene ontology (GO) terms associated with cytoplasmic translation and synapse biology (Figure 1f, Supplementary Table 1).

### Subtypes of cortical projection neurons show functionally distinct GC transcriptomes, which are enriched for transcripts associated with neurodevelopmental and neuropsychiatric disorders

We next asked whether and to what extent GC transcriptomes differ between subtypes. To optimize rigor, we excluded genes that significantly differ between the unpurified GCFs of the two subtypes, thus retaining 509 comparable genes. We find that 99 (16%) are enriched in CPN GCs and 185 (31%) are enriched in CThPN GCs (Figure 2a, Supplementary Table 2). GC-localized transcriptomes might arise from a combination of transcriptional distinctions between the subtypes of parent somata, or by subtype-specific differences in RNA trafficking and stability. Intriguingly, we identify that, while indeed GC-localized abundances of many genes are transcription-associated, a large proportion of GC-distinct genes appear to be driven by differences in RNA trafficking and stability (Figure 2 b-d). Some genes can be classified as transcriptionally regulated (“transcription”, class VI/VII), while others are clearly linked to trafficking phenotypes (“trafficking”, class IV/V and “trafficking inverted”, class VIII/IX). For these trafficking-related classes IV/V and VIII/IX, there are clear distinctions in GC transcriptomes between subtypes. In contrast, these trafficking-related classes do not show differences between transcriptomes in somata. Many neurodevelopmental and neuropsychiatric disorders, e.g. autism spectrum disorder (ASD) and schizophrenia, respectively, are linked with aberrant cortical circuit formation. Recent efforts by several groups have identified human genetic variants associated with these and other disorders, e.g. bipolar disorder. We cross-referenced our subcellular data with disease-associated gene sets, and identify highly significant enrichment, in both GCs and somata of both sub-types, for ASD-associated genes as defined by both coding variants enriched in whole exome sequencing^72^ and curated by SFARI^73^. We also identify significant enrichment for schizophrenia-associated genes^74,75^ in CThPN soma and GC transcriptomes (Figure 2e, Supplementary Table 3).

**Figure 2:**
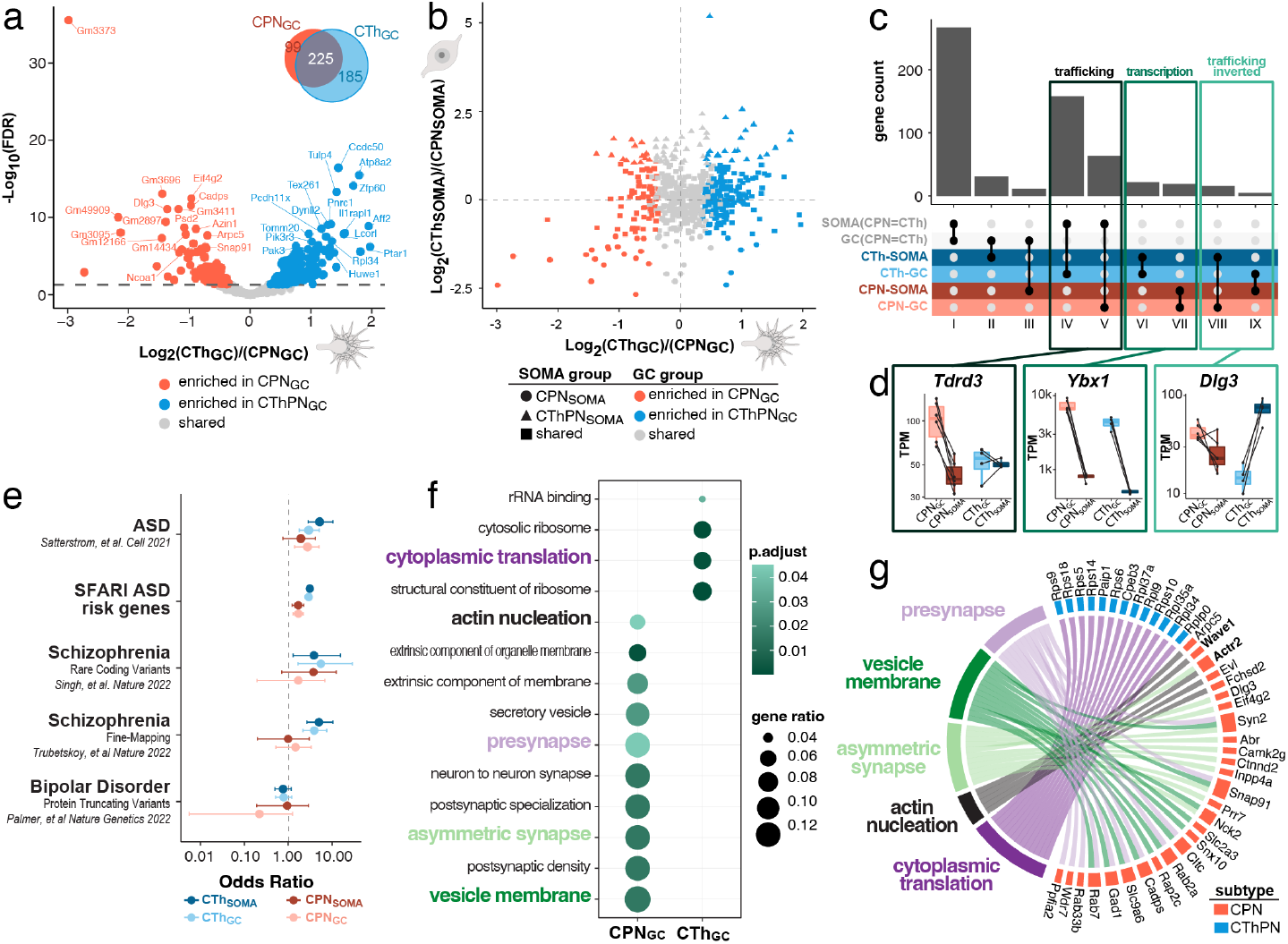
Distinct subtype-specific, GC-specific, local transcriptomes of CPN and CThPN are enriched with potential regulators of distinct axon guidance decisions. **(a)** Volcano plot indicating subtype-specific, GC-localized transcripts as colored dots (FDR. < 0.05). Venn diagram shows distribution of shared and distinct GC-specific molecular machinery between subtypes. **(b)** Transcriptomic differences between CPN and CThPN across subcellular compartments (somata *vs*. GCs). **(c)** Comparison of subtype-specific, GC-localized transcriptomes reveals molecular machinery that is shared between CPN and CThPN (classes I-III) and distinct between subtypes (IV-IX). Importantly, differences between GCs of distinct subtype are often not captured by comparing subtype-specific somata alone; we denote these as “trafficking” distinctions between subtypes (classes IV, V, VIII, IX), as opposed to those driven by “transcription” (classes VI, VII). **(d)** Normalized transcript abundances of exemplar genes for classes IV/V (“trafficking”, *Tdrd3*), classes VI/VII (“transcription”, *Ybx1*), and classes VIII/IX (“trafficking inverted”, *Dlg3*), across subcellular compartments and subtypes. **(e)** ASD and schizophrenia disease-associated genes are enriched (95% confidence interval) in soma- and GC-enriched transcriptomes. Both CPN and CThPN soma-localized and GC-localized transcriptomes show enrichment for ASD-associated genes, based on comparison with two independent data sets. Cross-referencing with two independent GWAS studies for schizophrenia additionally revealed that CThPN soma-and GC-localized transcriptomes are also enriched for schizophrenia-associated genes. There is no enrichment detected with genes associated with bipolar disorder in either CPN or CThPN transcriptomes, serving as a neuropsychiatric disease control, highlighting specificity in ASD and schizophrenia. **(f)** GO term enrichment in the GC-localized, subtype-distinct genes of each subtype. Color indicates -log10(P-value). **(g)** Subset of GO terms significantly enriched in either CPN or CThPN GCs connected with genes in GCs that are most frequently represented by those GO terms. Color of block proximal to each gene name denotes its subtype enrichment at P3. Bolded genes are core components/interactors of the WRC (Fig. 3).

We next investigated whether and which functionally coordinated groups of genes are subtype-specifically present in GCs. We identify that GO terms associated with ribosome biology and cytoplasmic translation are highly enriched with CThPN_GC_, whereas CPN_GC_ display significant enrichment for terms of actin nucleation and synaptic biology, likely reflecting differential developmental states and the corresponding stage-specific loads of RNA for local translation for the two subtypes (Figure 2f and Supplementary Table 4). We analyzed which differentially abundant genes contribute most significantly to the enriched GO terms associated with these categories. Intriguingly, we identify a number of core components and associated regulatory elements of the WAVE regulatory complex (WRC), e.g. *Wave1, Wave2, Actr2*, and *Abi1*, as GC-localized and, in some cases, subtype-distinct at P3 (Figure 2g).

### Transcripts of multiple WRC components and interacting receptors are enriched in GCs, some in a subtype-specific manner

The WRC is a pentameric complex consisting of WAVE1 (or its paralogs WAVE2 or WAVE3), CYFIP1 (or its paralog CYFIP2), HEM2 (or its paralog HEM1), ABI2 (or its paralogs ABI1 or ABI3) and BRICK. In its basal state, the WRC is autoinhibited in the cytosol, and gets recruited to the plasma membrane and activated, through cooperative action of GTPases (such as RAC1, ARF1, or ARF6), phospholipids (such as PIP3), kinases (such as ABL, SRC, and CDK5), and WRC interacting receptor sequence (WIRS) containing membrane receptors. Upon recruitment to the plasma membrane and activation, the activating subdomain WCA is released, which in turn stimulates the ARP2/3 complex to polymerize actin, thus mediating actin nucleation and branching (Figure 3a, schematic adapted from^1^).

**Figure 3:**
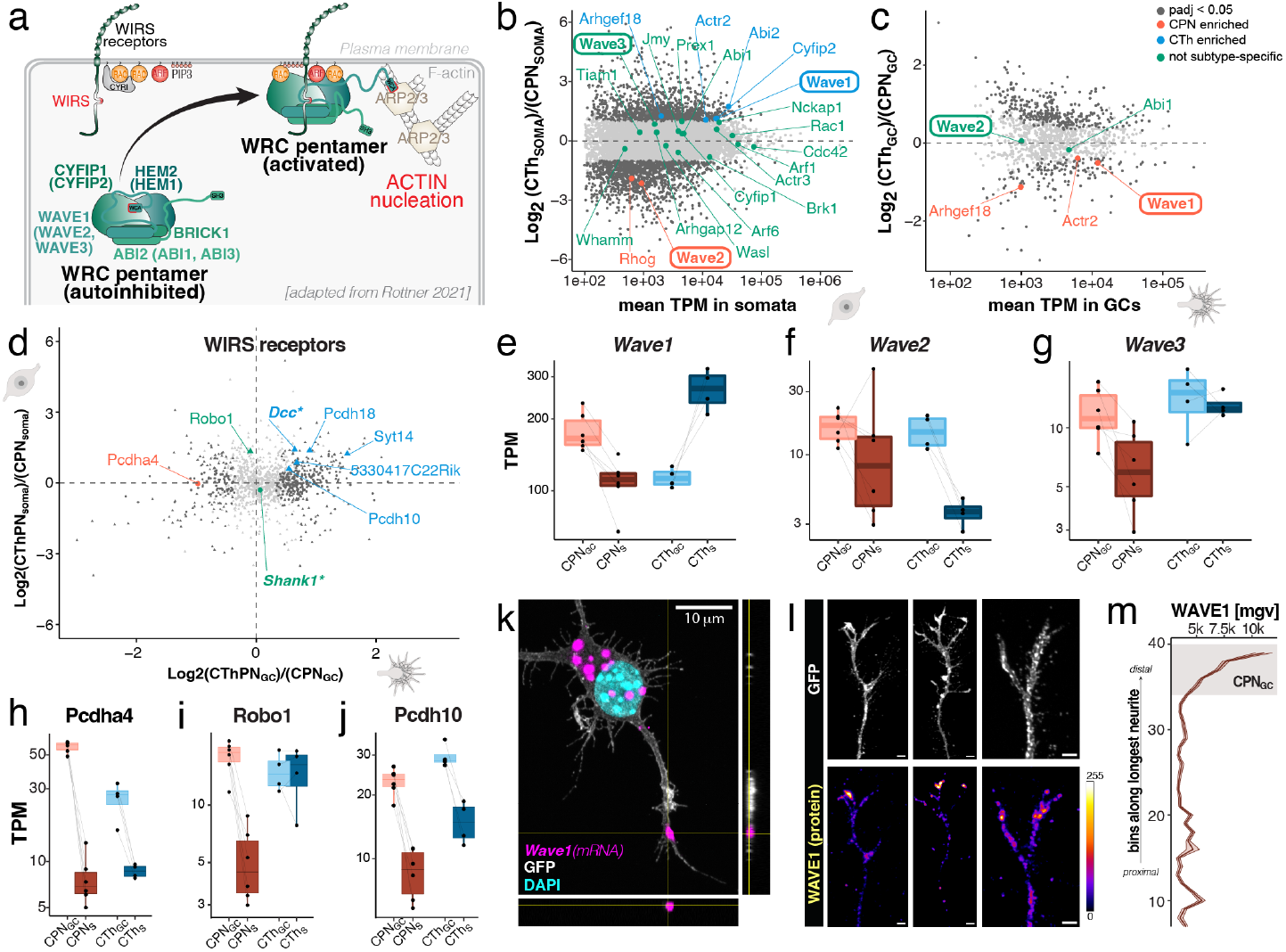
Core components and regulatory elements of the WAVE regulatory complex are subcellularly enriched in GCs of CPN and CThPN, suggesting local assembly and subtype-specificity of the machinery regulating actin branching. **(a)** Schematic of the composition, regulation, and function of the WRC (also termed SCAR complex), which is activated upon association with WRC-interacting regulatory sequence (WIRS)- containing receptors and is important for actin nucleation and branching. Adapted from Rottner 2021^1^. (b) WRC core genes and regulators: transcript abundances and subtype-specific expression in CPN and CThPN somata. **(c)** WRC core genes and regulators: transcript abundances and subtype-specific expression in CPN and CThPN GCs. **(d)** Localization plot showing subtype-specific localization of transcripts encoding high confidence WIRS-containing proteins. **(e-g)** Normalized transcript abundances of *Wave1, Wave2*, and *Wave3*, indicating their subtype-specificity and subcellular transcript localization. **(h-j)** Normalized transcript abundances of select WIRS-containing transcripts, showing their subtype-specificity and subcellular localization. **(k)** Maximum intensity projection (10 μm total z depth) of representative RNAscope confocal images of *Wave1* in GFP-positive cultured CPN. Localization of the magenta *Wave1*-positive puncta in the tip of the neurite is confirmed by respective orthogonal z-axis views along the sides. **(l)** Maximum intensity projection (10 μm total z depth) of representative immunocytochemistry for GFP (top panels) and WAVE1 (bottom panels) in GCs of cultured CPN. WAVE1 protein is detected as focal puncta at the tip of CPN GCs. **(m)** Quantification of WAVE1 immunocytochemical signal intensity (mean grey value, mgv ± sem, n=11) in the longest neurite of cultured CPN (n = 11), from proximal (bottom) to distal (top) end of the neurite. Grey box indicates localization of GC as assessed by parallel immunolabeling for actin.

We identify that both CPN and CThPN somata express all the necessary core components of the WRC (though *Abi3* and *Hem1* are detected at very low abundance), along with several secondary regulatory WRC interactors (Figure 3b and Supplementary Figure 5a-c). Investigation of GC-localized transcriptomes identifies only a subset of WRC-associated transcripts as locally enriched, some of them with intriguing, subtype-specific differential localization across neuronal subcompartments (Figure 3c, e-g and Supplementary Figure 4a/b/e). For six members of the Wiskott-Aldrich syndrome protein (WASP) family, i.e. the three *Wave* paralogs (*Wave1, Wave2*, and *Wave3*), *N-Wasp, Jmy*, and *Whamm*, expression is detected in both CPN and CThPN somata. Notably however, within this protein family, transcripts of only the three *Wave* paralogs (*Wave1, Wave2*, and *Wave3*) display differential subcellular localization between soma and GC compartments (Figure 3e-g and Supplementary Figure 5d). Of the three *Wave* paralogs, *Wave1* displays the highest expression levels and the largest difference in subcellular localization between subtypes (Figure 3e). We confirmed the presence of *Wave1* transcripts in neurites of cultured CPN (Figure 3k and Supplementary Figure 7), and enrichment of the WAVE1 protein product toward the distal end of the longest neurite of cultured CPN (Figure 3l & m).

The WRC is particularly relevant to axon guidance and circuit formation because it links extracellular cues sensed by transmembrane receptors to actin cytoskeletal regulation via WIRS-containing proteins. Transcripts encoding WIRS-containing proteins are differentially localized to GCs of both CPN or CThPN (Figure 3d, h-j and Supplementary Figure 6), though there is likely modest subtype-specific difference in localization of WRC-associated transcripts to GCs. Intriguingly, many of the WIRS-containing transmembrane receptors are reportedly associated with multiple neurodevelopmental disorders (e.g. *Dcc*^75,76^; *Shank1*^77^).

### UTRs of GC mRNAs are longer and enriched for translationally repressive cytoplasmic polyadenylation elements (CPEs)

We next investigated how mRNAs might be transported to and/or locally regulated in GCs. Since 3 ’UTRs are known to affect RNA trafficking and stability, we focused on comparison of distinct 3 ’UTRs between mRNAs. (Supplementary Figure 8).

We identified that 3 ’UTRs of GC-localized transcripts are significantly longer than those in somata, most significantly in CPN (Figure 4a). We scanned 3 ’UTRs identified as preferentially enriched in GCs for known motifs using the tool Transite^78^. We identified strikingly correlated motif enrichment in both CPN and CThPN GCs. Most strikingly, we identify U-rich sequences associated with the RBP families CPEB, RALY, and TIA1 (Figure 4b). To account for incomplete motif databases, we also searched for ungapped motifs *de novo* using STREME^79^. The top five logos identified by *de novo* motif analysis represent U-rich sequences, in agreement with results we obtained by motif scanning alone (Supplement Figure 8e).

**Figure 4:**
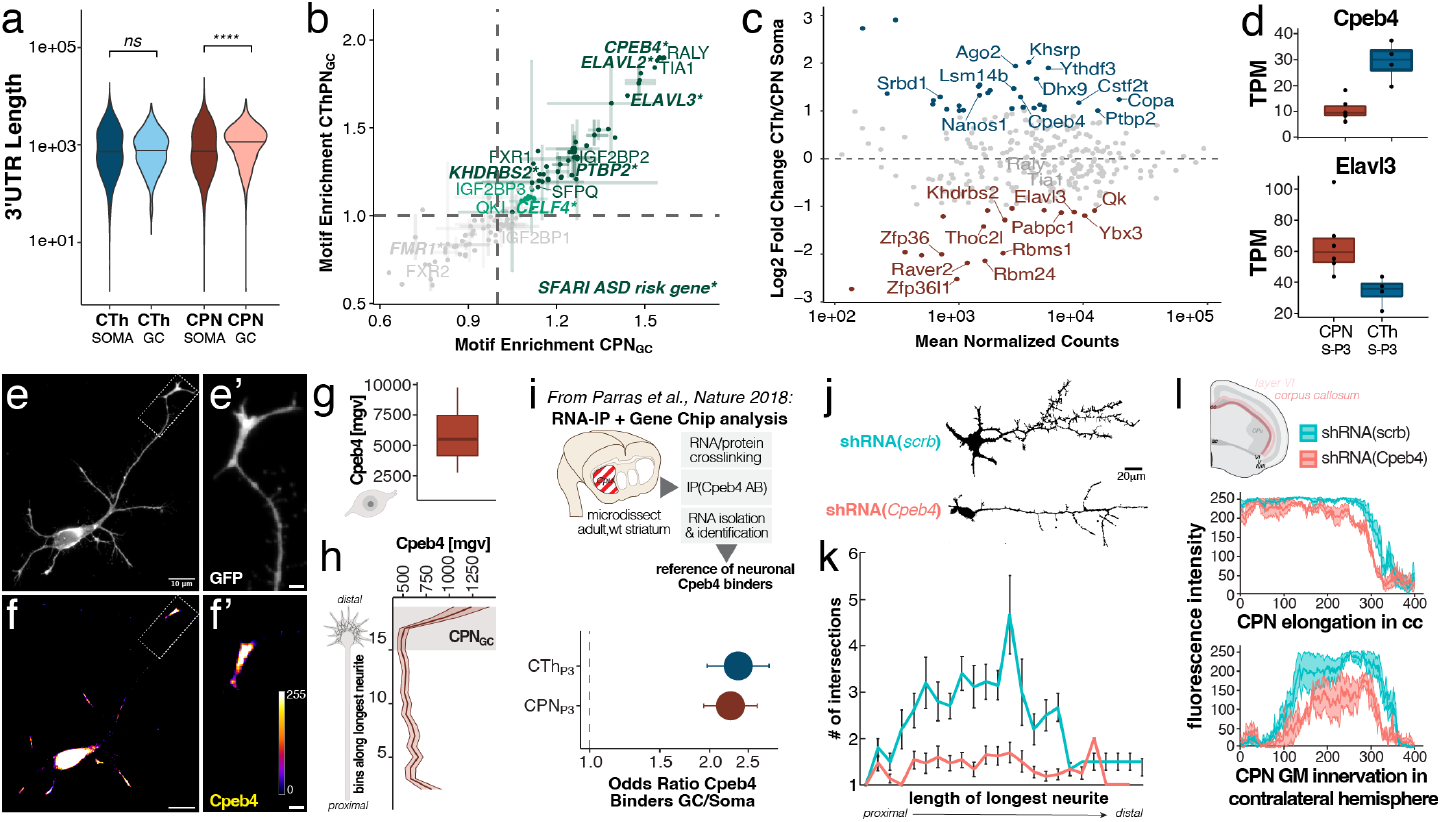
GC-localized mRNAs of both subtypes exhibit shared 3 ’UTR and motif characteristics, and are enriched for motifs associated with CPEB4. **(a)** 3 ’UTRs are on average longer in GCs than in somata. **(b)** Motif enrichment in GCs *vs*. somata is highly correlated between CPN and CThPN GCs; the most GC-enriched poly-U motifs are associated with CPEB4, RALY, and TIA1. **(c)** Comparison of CPN and CThPN soma-localized RNA expression of RBPs, annotated by GO term 0003729. Subtype-specific RBPs are highlighted in red (CPN) and blue (CThPN), FDR < 0.05 and fold change > 2. **(d)** RNA expression in CPN (red) and CThPN somata (blue) for *Cpeb4* and *Elavl4*. **(e & f)** Cultured mouse CPN expressing myristoylated GFP, with distal axon (longest neurite) magnified in (e ‘); immunolabeling for CPEB4 is shown in (f) with magnification showing distal axon enrichment of CPEB4 in (f ‘). **(g)** Quantification of mean gray value of Cpeb4 immunolabeling in somata of primary cultured CPN (n = 6). **(h)** Analysis of same gray value scale of CPEB4 immunolabeling intensity binned from proximal axon (bottom) to distal GC (top) for the longest neurite (n = 6). **(i)** CPEB4-bound mRNAs identified by RIP-seq^2^ are highly enriched in GCs of both subtypes as compared to their somata. **(j)** Cultured CPN fixed at 3 days *in vitro* (DIV3) display decreased axon branching complexity following *Cpeb4* shRNA-mediated knockdown. **(k)** Sholl analysis of branching complexity of longest neurite of cultured CPN, comparing neurons treated with *Cpeb4* shRNA (n = 15) or scrambled control shRNA (n = 15). Intersections (mean ± sem) with concentric spheres spaced every 10 μm around the cell body are quantified. **(l)** Constitutive shRNA knockdown of *Cpeb4* in CPN (red, n = 3) by *in utero* electroporation does not nonspecifically affect axons elongating into the corpus callosum, but it significantly reduces innervation of the contralateral gray matter compared to CPN treated with a scrambled shRNA control (scrb, teal, n = 4).

We next compared expression profiles of annotated RBPs between CPN and CThPN somata (Figure 4c). We find that all three families of RBPs with motifs enriched in GC transcripts are expressed by both subtypes (Figure 4c), with *Cpeb4* being more abundant in CThPN somata than in CPN somata at P3 (Figure 4c & d). Since CThPN and CPN are born several days apart during corticogenesis, and thus undergo hetereochronic development, differential abundances of RBP transcripts are likely due to differences both across subtype and across growth state.

### CPE-binding protein 4 (CPEB4) is enriched in distal neurites, suggesting function as a regulator of local translation

CPEBs are multifunctional RNA binding proteins that are reported to be involved in localization (CPEB1 and CPEB3^80-83^), translational regulation ^81,84-89^, and regulation of RNA stability ^90,91^. With respect to translation, CPEBs bind 3 ’UTRs of mRNAs in complex with other factors in an RNP that represses translation of the RNA until the CPEB is phosphorylated, which changes the RNP composition and ultimately enables translation (Supplementary Figure 9a). *Cpeb4* is the most highly expressed paralog of this family in CPN somata (Figure 4d and Supplementary Figure 9b). Its mis-splicing has been associated with ASD^2^, suggesting that it might function in regulating circuit formation in developing neocortex. Since we observed strong enrichment for CPEs in 3 ’UTRs of GC-localized mRNAs, we hypothesized that CPEB4 might perform subcompartment-specific regulatory function in developing cortical projection neurons.

Since motifs are a proxy for potential binding, we re-analyzed CPEB4 RNA-IP data published previously by Parras *et al*^2^. We find that experimentally identified RNAs bound by CPEB4 in the subcortical striatum are overrepresented in GC transcriptomes compared to soma transcriptomes (Figure 4i). Building on the result that transcripts with CPEs in their 3 ’UTRs are enriched in GCs, we further investigated potential involvement of CPEB4 in subcellular transcript regulation via immunocytochemistry for CPEB4 in primary cultured CPN. We identify that CPEB4 protein is enriched at the distal ends of neurites (Figure 4e-h). These data were further supported by western blot results comparing CPEB4 protein levels in forebrain overall, micro-dissected cortex, and CPN axon bundles micro-dissected from the corpus callosum. CPEB4 and its isoforms are detected in the axon compartment (Supplementary Figure 9c). We functionally investigated the effects of depletion of *Cpeb4* via shRNA-mediated knockdown in primary cultured CPN. We find that branching complexity of the longest neurite is severely reduced when *Cpeb4* is knocked down, compared to scrambled control (Figure 4j/k, Supplementary Figure 10a /b). We next assessed the effects of Cpeb4 knockdown in CPN in vivo, by selectively targeting CPN via E14.5 IUE of respective shRNA constructs (Supplementary Figure 10c/d), and saw a reduction in fibers innervating the contralateral grey matter, compared to scrambled control (Figure 4l).

It is known that the spatial characteristics of CPE placement in the 3 ’ UTR determines the proximity of potential binding partners (Supplementary Figure 9a) and subsequent effects of CPEB binding on translational output (Supplementary Figure 9d). These spatial characteristics include the polyadenylation signal, the hexamer binding site of cleavage and polyadenylation specificity factor (CPSF), and the distance between CPEs. Therefore, we profiled the location of CPEs in 3 ’UTRs of GC transcripts to investigate potential mechanisms by which CPEBs might regulate transcripts in GCs. The presence of a CPE spaced 6-25 nt from the CPSF hexamer binding site at the 3 ’ end of a transcript enables translational activation^92,93^, and the presence of a CPE in the 3 ’UTRs of several mRNAs has been associated with localization to neurites^94,95^.

We identify that the location of CPEs is within 120 nt of the most distal polyadenylation signal for >80% of GC transcript 3 ’UTRs. This suggests that CPEB4 binding in general can activate translation of these transcripts and might also facilitate transcript GC localization. In addition to activation, CPEBs can also mediate translational repression; shorter distances between CPEs results in translational repression. Such CPEB-mediated translational repression can facilitate cue-dependent regulation of local protein abundance (reviewed in^96^). In 3 ’UTRs of GC transcripts, CPEs are longer and more evenly distributed than in soma transcripts (Supplementary Figure 9e). Further, 3 ’UTRs in >80% of GC transcripts have CPE motifs spaced 10-50nt apart, a distance compatible with baseline translational repression (lower panel, Supplementary Figure 9e,^93^). We identify that a variety of versions of U-rich motifs are present in GC transcripts and soma-restricted transcripts of both subtypes, with a tendency for preferential usage of the most U-rich versions (TTTTTT, TTTTTG, (G/C/ATTTTT) in GC transcript 3 ’UTRs (Supplementary Figure 9f).

### GC transcripts exhibit dynamic, developmental stage-dependent changes in local machinery

CPN establish their subtype-specific circuitry in several distinct phases during late development. After exiting the ipsilateral cortical plate, their axons cross the midline via the corpus callosum in a highly fasciculated bundle and elongate through the more lateral subcortical white matter. Once axons have reached their appropriate homotopic target regions in the contralateral hemisphere, they defasciculate and innervate the cortical grey matter via branching and collateralization. They ultimately form synapses with appropriate target neurons (Figure 5a, Supplementary Figure 11, reviewed in^97^. During these distinct phases, overall gene expression shifts substantially, as CPN progress from elongating axon growth at P1 to grey matter innervation and initiation of synapse formation at P3. Gene set enrichment analysis of somata transcriptomes reflects this developmental progression (Supplementary Figure 12a).

**Figure 5:**
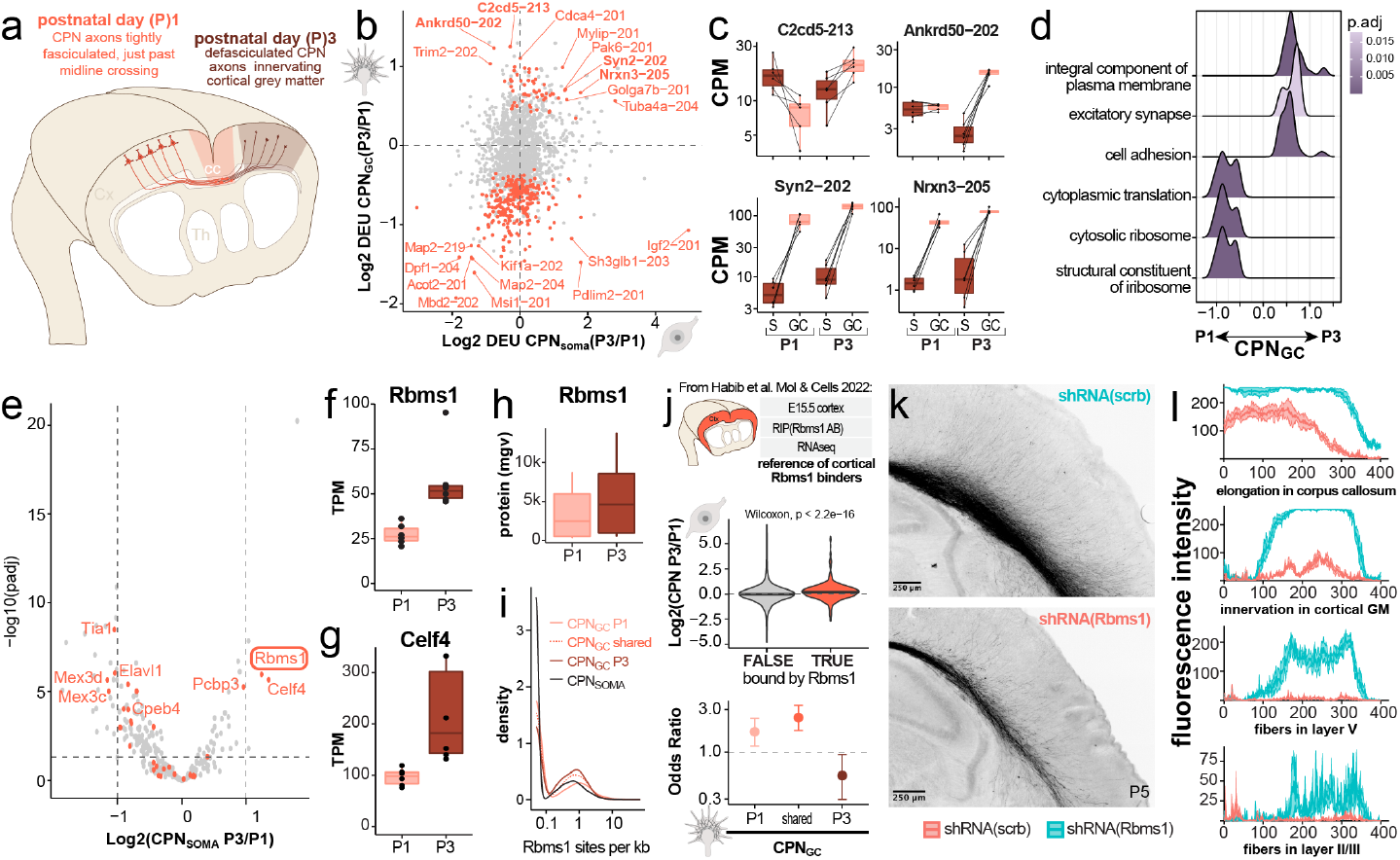
GC-localized transcripts change dynamically across developmental stages. **(a)** Schematic of CPN GC locations and growth state at P1 (fasciculated, just past midline crossing) and P3 (defasciculated, innervating contralateral gray matter containing target neurons). **(b)** Quadrant plot displaying changes in 3 ’UTR isoform usage (differential exon usage, DEU) from P1 to P3 in CPN somata (x-axis) and GCs (y-axis), respectively. The most significant changes in transcript isoform usage are found in GCs, resulting in a predominantly vertical distribution. **(c)** Boxplots highlighting examples of temporally shifting candidate transcript isoforms, displaying TPM across compartments (GCs pink, somata brown) and developmental stages (P1 vs. P3). **(d)** Gene set enrichment analysis comparing functional sets of transcripts changing from P1 to P3 in GCs reveals an increase in GC-local transcripts associated with cation transport and excitatory synapse formation. **(e)** Differential RNA expression comparing CPN somata at P3 *vs*. P1 of RBPs annotated by the GO term 0003729; orange dots indicate RBPs; grey dots indicate non-RBP transcripts. Labeled dots highlight genes that are associated with motifs enriched in GCs over somata (see Fig 4). **(f & g)** Boxplots highlighting RNA expression changes of *Rbms1* and *Celf4*, the two RBPs for which expression by CPN increases the most from P1 to P3, and which likely have GC-localized functions. **(h)** Mean gray value of RBMS1 protein labeling in somata of primary CPN plated at P1 (pink) or P3 (brown). **(i)** Motif scanning for RBMS1-associated motifs in 3 ’UTRs detected as GC-localized at P1, P3, or at both developmental stages (shared). Lighter and darker shades of red indicate the proportion of RBMS1-associated motifs detected in respective 3 ’-UTRs. **(j)** Cross-referencing of subcellular transcriptomes identified here with previously published RBMS1 RIP-seq data from homogenized embryonic cortex^21^ reveals higher abundances of RBMS1-bound transcripts at P3 *vs*. P1 in CPN somata (middle panel), but de-enrichment of RBMS1-bound transcripts in CPN GCs at P3 *vs*. P1 (lower panel). **(k)** Constitutive shRNA knockdown of *Rbms1* (red, n = 4) in CPN by *in utero* electroporation results in fewer axons elongating into the corpus callosum (*cc*) and innervating the contralateral gray matter when compared to CPN treated with a scrambled control shRNA (scrb, teal, n = 5), quantified in **(l)** as mean gray value (mgv, mean ± sem).

We investigated the dynamics of subcellular CPN transcriptomes by comparing their GC transcriptomes at P1 and P3. We identify increased 3 ’UTR length for P3 CPN_GC_ when compared to P1 CPN_GC_ (Supplementary Figure 12d). Assessment of functional categories of transcripts that dynamically change in CPN GCs through development (Figure 5b and Supplementary Figure 12b & c) reveal that gene sets associated with excitatory synapse formation, transmembrane cation transport, and cell adhesion are enriched in P3 CPN GCs over P1 CPN GCs (Figure 5d). For example, we identify GC-specific changes in transcript abundances of *C2cd5, Ankrd50, Syn2*, and *Nrxn3*, which are all involved in formation and maintenance of excitatory synapses (Figure 5c,^98-101^).

### Dynamic changes highlight potential regulatory roles for RBPs such as RBMS1

We investigated RBPs in CPN whose expression levels change over time, highlighting them as potential regulators locally mediating subcellular transcriptomic changes. We identified *Rbms1, Celf4*, and *Pcbp3* as increasing significantly over developmental time (Figure 5e-g). Integrating previously published microarray data^51^, *Rbms1* expression is found to peak at ∼P3, suggesting that it might likely function specifically in development of CPN at this crucial transition from elongation growth to collateralization/synapse formation (Supplementary Figure 13a). *Rbms1* paralogs *Rbms2* and *Rbms3* have very low transcript abundances at both P1 and P3 (Supplementary Figure 13b), suggesting that *Rbms1* is the dominant paralog in this system. RBMS1 protein product is present in the somata of primary cultured CPN at P1 and P3 (Figure 5h). Intriguingly, transcripts that are enriched in GCs have an increased number of Rbms1 motifs in their 3 ’ UTRs, an effect most pronounced for GC transcripts at developmental stage P1 (Figure 5i).

We analyzed the GO terms of CPN soma and CPN GC transcripts that contain RBMS1 motifs, revealing enrichment for terms associated with ion transport and membrane biology (Supplementary Figure 13c). We cross-referenced previously published data identifying putatively RBMS1-bound transcripts in developing cortex^21^ with our CPN-specific, subcellular data. Again intriguingly, transcripts bound by RBMS1 are more likely to be GC-localized at P1 compared with P3 (Figure 5j, lower panel). Further, RBMS1-bound transcripts display higher overall abundance in somata at P3 compared with P1 (Figure 5j, middle panel). Together, these dynamic data suggest that RBMS1 functions in part to stabilize transcripts.

Since RBMS1 has been previously reported to be important for CPN radial migration^21^, and for regulation of tissue invasion and metastasis in various forms of cancer^102-104^, we hypothesized that manipulation of *Rbms1* specifically in CPN *in vivo* might cause deficits in the ability of the manipulated CPN GCs to innervate gray matter on the contralateral hemisphere. We performed shRNA mediated knockdown of *Rbms1* via unilateral IUE at E14.5 (Supplementary Figure 13d-g). This results in pronounced reduction in the number of CPN axons crossing the corpus callosum, CPN axons innervating the contralateral grey matter, and CPN axon fiber density across all layers of contralateral cortex, compared to scrambled control (Figure 5k & l). Together, these results support an important and dynamic role for GC-localized RBMS1 in CPN contralateral axon connectivity and innervation, critical for circuit formation.

## Discussion

The work presented here has implications well beyond the specifics of brain circuitry development, maintenance, and function. Most differentiated cell types, whether highly polarized or not, contain multiple subcellular compartments with highly specialized local functions – e.g. apical vs. basal domains in epithelia; extending processes and sites of intercellular engagement in fibroblasts, oligodendroglia, and many immune populations. Neurons, and in particular cortical projection neurons, are extreme examples of highly polarized and subcellularly compartmentalized cells, with critical need to distribute function- and location-specific molecular machinery across vast spatial expanse. This subcellular localization enables axons, dendrites, GCs, and synapses to function and dynamically adapt in a semi-autonomous fashion far from the cell body and nucleus. Especially during development, when the distance between axon and respective parent soma is growing increasingly larger, GCs must be locally equipped with all the molecular machinery required to navigate selective guidance cues and to successfully establish subtype-specific circuitry. In the most extreme examples, peripheral dorsal root ganglion sensory neurons and select subtypes of cortical projection neurons, e.g. many subtypes of CPN^105^, corticostriatal PN^106^, and SCPN^107^, extend multiple separate axons that originate from the same parent soma, in parallel, to distinct target areas. In addition to the high complexity in subtypes of cortical projection neurons, cortical circuitry is additionally characterized by pronounced point-to-point connectivity, thereby enabling highly advanced sensory, motor, and cognitive function. The diversity and function-specificity of these selectively “wired” circuits, plus the critical nature of precise and subtle associative and cognitive function, makes cortical connectivity specifically vulnerable to effects of mis-wiring. Even subtle variations and/or abnormalities can lead to aberrant function– from mild to severe phenotypes of neurodevelopmental and neuropsychiatric disorders, e.g. autism spectrum disorders, intellectual disabilities, or schizophrenia.

In this work, we investigate *in vivo* GC and soma transcriptomes from two sharply distinct subtypes of cortical projection neurons over developmental time. CPN and CThPN, while highly related subtypes with regard to their pallial progenitor origin and general capabilities as cortical projection neurons, set up vastly different axonal trajectories over development, leading to dramatically distinct functions.

### Comparative transcriptomics of cell bodies and GCs, isolated in parallel from the developing mouse brain, enables investigation of subcellular molecular machinery

Our work extends upon studies that investigated local transcriptomes and translation in the context of formation^29,36,108-112^, maintenance^113-115^, and plasticity of synapses^116-119^. These studies have established the need for subcellularly distinct transcriptomes and local translation for pre- and post-synaptic function (reviewed in^120^).

In contrast, little is known about spatial segregation of molecular machinery in the context of development and circuit formation *in vivo*. A rare exception is the work of Shigoeka et al. employing TRAP to investigate axonal changes in translated RNAs over the time course of RGC circuit formation. They found that the RGC-axonal translatome is dynamically regulated during development, and that targets of mTORC1, FMRP, and APC display translational co-regulation in a stage-specific manner^121^. Our lab has combined biochemical fractionation of GCs, originally established by Karl Pfenninger and colleagues for forebrain GC isolation^122^, with subtype-specific fluorescent labeling and purification of GCs by newly developed fluorescent small particle sorting. This enables parallel investigation of soma-*vs*. GC-localized transcriptomes^7,50^. In the first application of these new experimental and analytic approaches, Poulopoulos*, Murphy* et al. provide a first global delineation of the P3 developing mouse CPN GC transcriptome and proteome. This work identified and validated by multiple standard methods preferential localization and function of 5 ‘TOP motif mTOR hyper-sensitive transcripts to the GC compartment during axonal projection *in vivo*^7^.

### GC-transcriptomes of different subtypes of cortical projection neurons are dynamically regulated and contain core, shared aspects, as well as context-specific machinery

Here we utilize the same set of experimental approaches to selectively investigate and map the GC *vs*. soma transcriptomes of CPN and CThPN across distinct stages of circuit formation. Within the GC-localized transcriptomes, we identify a core set of genes involved with cytoplasmic translation and synapse formation that are shared between subtypes. Additionally, we identify subtype- and stage-specific specializations of the transcriptomes, indicating and elucidating context-dependent regulation of mRNA content in GCs. As examples, we identify known and novel regulators of GC Rac1-associated signal transduction, actin cytoskeletal regulators including elements of the WRC, and genes involved in early selection and development of synapses. Consistent with our lab ’s prior work, we identify dynamic expression of ribosomal mRNAs, which decrease over time in the GC compartments of both subtypes. This is consistent with the reduced required load of protein synthesis in later development that has been reported previously^121^, as GCs transition from rapid growth in fasciculated bundles to slowed growth and innervation of gray matter with subsequent synapse formation. We also identify an increase in gene sets associated with membrane biology, cell adhesion, and early synaptogenesis, as well as select candidates that previously have been associated with neuronal migration and tumor invasiveness (e.g. *Dcx*^123^), likely reflecting a molecular shift important for target tissue innervation, e.g. extension through complex interstitial spaces.

### Directed transport as well as tight regulation of RNA stability and translation are crucial for establishing distinct subcellular transcriptomes

Intriguingly, we identify– both by comparing subtypes of projection neurons and by investigating across distinct developmental stages– that the soma-localized transcriptome is not fully predictive of the GC transcriptome at the same time. Several known levels of regulation are likely to contribute to this striking result. These include long-range directed transport^11,13-20,95,124,125^ and RNA stability^23,24^ (reviewed in^25^), which influence both the rate at which RNAs arrive at GCs and the duration of time they are available within the GC-localized transcriptome, respectively. We identify multiple characteristics of GC-localized RNAs that likely contribute to subcellular localization and translational regulation in the context of developmental dynamics. First, we identify that the length of 3 ’UTRs positively correlates with developmental stage, consistent with previous observations^126-128^. The lengthened 3 ’UTRs represent expanded spatial extent of associated sequences, which likely facilitate tighter control over subcellular localization, stability, and local translation by a suite of RNA binding proteins and miRNAs. Second, we identify a set of poly-U/CPE motifs enriched in 3 ’UTR sequences of GC transcripts, which are known to be regulated by several RBPs, including CPEB4.

### CPEB4 as potential integrative player in translational repression of GC-trafficked transcripts; dysregulation results in aberrant circuit formation and might be linked to ASD

We identify that Cpeb4 mRNA and CPEB4 protein are enriched at distal neurites, emphasizing its likely role in local regulation of transcript abundance and translational output in the GC compartment. CPEB family members have been shown to establish polarity of RNAs in oocytes (reviewed in^92^) as well as to facilitate neurite-directed mRNA transport in neurons^82,84,87^. CPEBs also modulate local translation by transducing a phosphorylation cue into increased transcript polyadenylation, with subsequent increase in translation initiation^81,85,86,88,89^. CPEB4 specifically has been shown to disproportionately bind identified Simons Foundation “SFARI” ASD risk genes, and misexpression of non-neuronal isoforms of *Cpeb4* in idiopathic ASD brains leads to decreased poly-A tail length of SFARI risk genes^2^. This is in line with the observation that SFARI risk genes are in general less efficiently translated compared to the rest of the transcriptome^129^. We find that the overall composition of CPE motifs within the 3 ’UTR of GC-enriched transcripts seem to favor baseline translational repression, but that the general features enabling CPEB4-mediated translational activation are also present in these UTRs. This suggests the hypothesis that CPEB4 and related RBPs might likely ensure overall translational repression of transcripts being trafficked to and stored at projection neuron GCs, locally poising them to be translationally activated in a likely coordinated fashion upon appropriate guidance cue signaling, via phosphorylation of CPEB4. Such translational repression in long-range RNA transport has been observed previously^130^; intriguingly, TIA1, which also binds poly-U motifs, has been related to stress granules^131^. Taken together, the combination of prior work with our results that ASD-associated genes are enriched in GCs leads to the intriguing hypothesis that CPEB4-mediated, cue-dependent translation specifically in developing GCs is compromised in ASD, and that this leads to developmental GC navigational abnormalities and aberrant circuit formation.

### RBMS1 regulated stability of GC-transcripts dynamically across developmental stages, and is likely a key player for successful tissue invasion

Our results further highlight that GC RNA regulation is dynamic, modulating transport and local stability of transcripts in a developmental stage-specific manner. As an example, we identify that the RBP *Rbms1* is upregulated as CPN GCs enter the specific phase of grey matter innervation and subsequent synapse formation. Intriguingly, we identify a decrease in RBMS1-bound RNAs in GCs from P1 to P3, while abundances of the same transcripts increase in somata. AU-rich sequences have been described as strong axon-targeting elements in UTRs^95^, raising the question of whether RBMS1 might act not only by stabilizing distinct sets of transcripts at specific times, as previously described^21,103,104^, but also by sequestering these transcripts to the cell body. Since RBMS1 is known to regulate stability of mRNAs in the context of metastasis and tissue invasion by several cancer types^102-104^, and since we find that knockdown of this RBP specifically in CPN results in decreased competency of projections to innervate gray matter and establish circuitry, we hypothesize that RBMS1 is a key regulator involved in tissue invasion by CPN.

## Conclusions

Understanding the neuronal subtype-specific composition of subcellular molecular machinery of axon and GC compartments, and its dynamic local regulation across subtypes and stages of circuit development, enables mechanistic dissection of regulatory switches between crucial growth states, whose mis-regulation results in aberrant circuit formation. Our work highlights the importance of investigating spatial distribution of subtype- and circuit-specific RNA and protein molecular machinery, and its likely location-dependent function, across sub-cellular compartments *in vivo*. Investigation *in vivo* enables access to and delineation of developmental stages and physiological complexity. This biology is missed both *in vitro* and when restricting investigation to somata *in vivo*. Increasingly deep mechanistic knowledge of these local regulatory mechanisms will contribute importantly to understanding connections between diverse and precise circuit development, disease risk genes, and neurodevelopmental and neuropsychiatric disorders.

## Acknowledgements

We thank J. LaVecchio and N. Kheradmand of HSCRB-HSCI Flow Cytometry Core; the Bauer RNA sequencing core; Harvard Center for Biological Imaging for infrastructure and support; and T. Sackton for statistical advice. This work was supported by grants to J.D.M.: Allen Distinguished Investigator Award ADI 11855 from the Paul G. Allen Frontiers Group; NIH Pioneer Award DP1 NS106665; and the Max and Anne Wien Professor of Life Sciences fund; with additional infrastructure support from NIH grants NS045523 and NS104055.

A.K.E. was supported by a Jean-Jacques et Felicia Lopez-Loreta Foundation Award and a Swiss National Science Foundation Postdoctoral Fellowship. P.V. was partially supported by NSF GRFP 280932, The NSF-Simons Center for Mathematical and Statistical Analysis of Biology at Harvard award number #1764269, and NIH Training Grant T32 GM007306-43. J.J.H. was partially supported by NIH NRSA F31 NS103262 and NIH Training Grant T32 GM007226. This paper was typeset with the bioRxiv word template by @Chrelli: www.github.com/chrelli/bioRxiv-word-template

## Author contributions

J.J.H. and J.D.M initiated the study. J.J.H. and A.K.E. constructed mouse lines.

T.A.A. and P.V. designed constructs. P.V. and J.J.H. performed *in utero* electroporations. A.K.E. and J.J.H. carried out GC preps and GC and soma sorts. Y.I. performed soma preps. P.V., A.K.E., and D.N. designed and executed cellular and histological validation studies. P.V. performed all bioinformatic analyses. P.V., A.K.E., and J.D.M. interpreted the data and iteratively designed further analyses and experiments. P.V. and A.K.E. drafted the manuscript. P.V., A.K.E., and J.D.M. edited the manuscript with input from the other authors.

## Materials and Methods

### Mice

Experiments involving mice were performed with approval of the Harvard University Institutional Animal Care and Use Committee. All CPN were isolated and purified from wildtype CD1 mouse pups of both sexes (Charles River Laboratories), enabling subtype-specific expression of fluorescent proteins using unilateral *in utero* electroporation at embryonic day 14.5 (E14.5), targeting CPN of cortical layers II/III. Experiments isolating and purifying CThPN were performed on BAC transgenic Ntsr1::Cre mice (MMRRC, 030648-UCD) backcrossed to CD1 for >10 generations. Ntsr1::Cre mice were bred with CD1 Emx1::FlpO+/+/Ai65(RCFL-tdT) mice, resulting in highly selective expression of tdTomato in CThPN of cortical layer VI, while eliminating off-target labeling of rare neurons in multiple midbrain structures (labeled in simpler non-intersectional crosses of Ntsr1::Cre with TdTomflox/flox). For maximal consistency, breeding pairs were co-caged for <12 hours, and postnatal times were defined as “postnatal day” (P) with most pups being born at *P*0.

### *In utero* electroporation

Subtype-specific labeling of CPN was achieved using unilateral *in utero* electroporation of a genetic construct of interest at E14.5, as previously described^132^. Briefly, a pregnant dam was anesthetized using 1.5-2% isoflurane, and prepared for surgery by applying eye ointment (#2444062, Systane) and administering buprenorphine (#60969, Par Pharmaceutical). The hair on the lower abdomen was removed using hair removal cream, and the skin was locally disinfected using ethanol. An incision was made through skin and abdominal musculature, and the uterine horns were exposed, gently lifted out of the intraperitoneal cavity, and placed on a sterile gauze pad where they were continuously kept moist using sterile, pre-warmed PBS. Embryos were individually positioned and injected unilaterally into one lateral ventricle (roughly alternating left and right) with 0.7-1 μl DNA solution (5 μg/μl), supplemented with Fast Green FCF (#AAA16520-06, Thermo Scientific Chemicals) for visualization, using a pulled and beveled glass micropipette (#22-260-943, Fisher). DNA was electroporated into neural progenitors lining the ventricles by applying current to the injected side of the cortex (5 pulses, 50 ms ON/950 ms OFF, 34 V), targeting lateral sensorimotor areas of the cortical plate. Following electroporation, embryos within their uterine horn were gently placed back into the intraperitoneal cavity, and the wound was closed using sutures (#39010, Covetrus). The dam was allowed to fully recover from anesthesia on a regulated heat mat before being returned to the mouse housing room. Postoperative care following surgery included daily checks of the mouse ’s overall well-being and its wound healing, and administration of buprenorphine (#60969, Par Pharmaceutical) every 12 hours for the first 3 days post-surgery. After term birth, mouse pups were screened for unilateral cortical fluorescence using a fluorescence stereoscope; successfully electroporated mice were marked with a tail clip.

### DNA constructs

Fluorescent labeling: We modified addgene plasmid #26771 to incorporate myr-tdTomato-2A-H2B-GFP (pCAG-myr-tdTomato-2A-H2B-GFP). This enabled neuronal projections in red and the nuclei in green. For culture experiments we instead incorporated myrmClover into addgene plasmid # 26771 (pCAG-myrmClover).

shRNA: MISSION shRNA plasmids (Sigma-Aldrich; see Table 1) at 2.5μg/μL were mixed with pCAG-myr-mClover or pCAG-myr-TdTomato plasmids at 2.5μg/μL and electroporated at E14.5 to target CPN, or at E11.5 to target CThPN.

**Table 1:** shRNA constructs.

| shRNA catalog number (Sigma) | target |
| --- | --- |
| TRCN0000096783 | ms-Rbms1 |
| TRCN0000096781 | ms-Rbms1 |
| TRCN0000197915 | ms-Cpeb4 |
| TRCN0000435567 | ms-Cpeb4 |
| SHC016 | non-target control (scrb) |

Over-expression constructs: Cpeb4 isoforms were cloned out of oligo-dT primed cDNA using gene-specific primers that included a 5 ’ HA tag sequence. These were inserted into a pCAG-[MCS]-IRES-GFP construct (CBIG), generated by C. Lois. Constructs were electroporated at 1μg/μL in conjunction with pCAG-myr-TdTomato at E14.5 to target CPN or at E11.5 to target CThPN.

DNA constructs were prepared using the endotoxin-free Maxiprep kit (Qiagen, #12362), following the manufacturer ’s instructions. Quality was verified using restriction enzyme digests, and by Sanger sequencing (Eton Biosciences).

### Assessment of shRNA knockdown efficiency

*In vitro*: Constructs were transfected into N2a cells using Lipofectamine 3000 (#L3000008, Thermo), following the manufacturer ’s instructions. For assessment via ICC, cells were re-plated on PDL-coated glass coverslips for an additional 24h, then fixed and stained according to the immunocytochemistry approaches detailed below. For knockdown assessment via qPCR, cells were sorted after 48 hours.

*In vivo*: Constructs were mixed with fluorescent reporter constructs (pCAG-myr-tdTomato-2A-H2B-GFP) and *in utero* electroporated at E14.5, targeting CPN of cortical layers II/III. On the day of birth (P0), fluorescence-positive neurons were purified using soma preparation and FACS approaches described below.

For both *in vivo* and *in vitro* applications, knockdown efficiency was calculated via qPCR: RNA was extracted with the Zymo Direct-Zol RNA microprep kit (#R2062, Zymo); cDNA synthesis was performed with the SuperScript IV First Strand Synthesis System using oligo-dT priming (#18091050, Thermo); and qPCR was performed with PowerUp SYBR Green Master Mix (Thermo, #A25918) using primers listed in table 2.

**Table 2:**
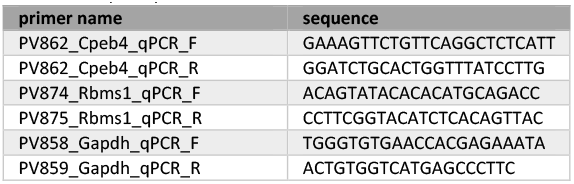
qPCR primers.

### GC fractionation & fluorescent small particale sorting (FSPS)

Isolation of GC fractions was performed using modifications of methods originally described by Pfenninger and colleagues^122^; these modifications are described in detail elsewhere^50^. Briefly, 8-10 brains per sample of mouse pups at selected developmental stages P1 or P3 were rapidly removed from the skull, and micro-dissected on ice in 0.32 M sucrose buffer supplemented with 4mM HEPES, HALT protease and phosphatase inhibitors (#78442, Thermo), and 1U/mL RNAse inhibitor (#N2515, Promega). CPN GCs at P1 were collected from the corpus callosum in the ipsilateral cortical hemisphere, just before midline crossing, where the main tract is located at this stage. Micro-dissections were performed on coronal sections under a fluorescence stereomicroscope at high magnification, carefully excluding any labeled somata from the micro-dissected area. CPN GCs at P3 were collected from the contralateral corpus callosum and cortical hemisphere. CThPN GCs at P1 and P3 were collected from the micro-dissected ipsilateral thalamus. For both subtypes of cortical projection neurons, micro-dissected tissue was supplemented with non-labeled forebrain tissue obtained from P3 mouse pups to achieve total tissue mass of 0.4 – 0.5 g.

Tissue was homogenized using a variable speed motor drive homogenizer (Glas-col, #099C K54) with 11 strokes at 900 rpm in a glass-Teflon potter. Post-nuclear homogenate (PNH) was obtained as supernatant after centrifugation at 1,600 g for 15min. PNH (in 0.32 M sucrose) was layered onto a 0.83 M/2.5 M sucrose cushion, and spun at 4°C in a fixed vertical rotor (VTi50, Beckman) at 242,000 g for 47 min to achieve subcellular fractionation.

Following ultracentrifugation, the GC fraction (GCF) was extracted from the 0.32 M – 0.83 M interface, and pre-labeled, subtype-specific, fluorescent GCs were purified using fluorescent small particle sorting (FSPS) on a customized SORP FACSAriaII (BD Instruments) equipped with a 70 μm nozzle running at 70 psi and a 0.1 μm filter for the sheath fluid to remove potential dust particles. The instrument is additionally customized to include an extra photomultiplier tube (PMT) to detect forward scatter signal, and a 300 MW 488 nm laser with reduced beam height (6±3 μm), as well as a custom lens assembly with noise-reducing filter and pico-motor focus. Scatter measurements were based on signal peak height, and plotted in log mode. To optimally adjust drop delay settings and check alignment of optics, sub-micron polystyrene size-standard beads (BDSORP beads, Spherotech, #NPPS-4K) were used to calibrate before each sorting experiment. Due to high sucrose content with high viscosity, GCFs were diluted in pre-cooled PBS before FSPS, and particles were gated for size and aggregation (based on SSC over FSC-PMT), and for fluorescence intensity (GFP or tdTom over FSC-PMT). A minimum of 500,000 to 1 million subtype-specific, fluorescently labeled GC particles were collected in RLT (Allprep DNA/RNA Micro Kit #80284, Qiagen) supplemented with 10% b-Mercaptoethanol.

### Western blot analysis and quantification

Western blots were performed using standard tris-glycine SDS-PAGE protocols. Samples had normalized protein concentration assessed using a Bradford assay (#23246, Thermo). Resolved proteins were electroblotted onto PVDF membranes using semi-dry transfer, blocked for 30 min in TBS supplemented with 0.1% Tween-20 and 5% skim milk, and incubated with primary antibodies (GAP43 #MAB347 Millipore; Actb #A5441 Sigma; Gm130 #610823 BD Biosciences; MAP2 #M1406 Sigma; WAVE1 #ab272916 abcam; WAVE2 #3659T Cell Signaling Technologies; WAVE3 #2806S Cell Signaling Technologies; Cpeb4 #ab224162 abcam; Rbms1 #ab150353 abcam; Tau #AF3494 RR&D) in TBS supplemented with 0.1% Tween-20 and 3% BSA over night at 4°C. The next day, the membrane was washed three times in TBS supplemented with 0.1% Tween-20, incubated for 2 hours at room temperature with secondary antibodies that were isotype-specific, HRP-conjugated, and cross-absorbed (Life Technologies), then washed again three times with TBS supplemented with 0.1% Tween-20. Immunoreactive bands were visualized through detection of chemiluminescence using a CCD camera imager (FluoroChemM, Protein Simple). Quantification of signal intensity in the various bands was performed using the standard Fiji software package.

### Hydrolysis (RNase or protease) protection assay on GCF

Membrane sealing and thus protection of GC-specific molecular contents was assessed using either an RNase or protease protection assay, as described in detail elsewhere^50^. In brief, isolated GCF samples were treated in parallel with either nothing (control; ctrl), 0.1% TritonX-100 detergent (D-ctrl), RNAse/protease only (R/P-ctrl), or a combination of 0.1% TritonX-100 and either RNase or protease (D+R/P) and incubated for 2 hours at 4°C. For RNase protection assays, samples were processed for RNA extraction, and quality of extracted RNA was assessed using an Agilent Tapestation 2100 with a High Sensitivity RNA kit, using the manufacturer ’s standard protocol. For protease protection assays, samples were processed for western blotting as described above, and probed with primary antibodies against the subcellular markers GAP43 (#MAB347 Millipore), Actb (#A5441 Sigma), Gm130 (#610823 BD Biosciences), and MAP2 (#M1406 Sigma), followed by secondary antibodies that were isotype-specific, HRP-conjugated, and cross-absorbed (Life Technologies).

### Subtype-specific soma isolation and purification by FACS

Dissociation of neuronal somata followed by subtype-specific purification by FACS was performed as previously described^50,71^. In short, mouse pups of the selected developmental age (P1 or P3) – either electroporated at E14.5 to label CPN or labeled by transgenes Ntsr1+/-, EmxFlpO+/-, Ai64+/-to label CThPN – were deeply anesthetized in ice, decapitated, and brains were extracted from the skull. Fluorescently labeled neurons in layer II/III (CPN) or layer VI (CThPN) were quickly micro-dissected in ice-cold HBSS under a fluorescence stereoscope, and transferred to a falcon tube containing dissociation solution. Tissue pieces were washed twice with 3-5 ml DS, then enzymatically dissociated by incubating twice with 5 ml enzyme solution, supplemented with DNAseI) for 15 min per incubation. ES was removed, and the tissue was washed twice with washing solution, followed by mechanical dissociation using gentle trituration with a fire polished Pasteur pipet. Dissociated CPN or CThPN were washed in 10 ml WS, and pelleted at 80g for 5 min at 4°C. The supernatant was removed, and neurons were resuspended in 500 μl WS using a fire polished Pasteur pipet. Subtype-specific, labeled CPN or CThPN somata were sorted into RLT buffer (Qiagen) supplemented with 10% b-Mercaptoethanol using a customized SORP FACSAriaII (BD Instruments) equipped with a 300 MW 488 nm laser set to 100 MW with large beam height and an 85 μm nozzle run at 45 psi.

### RNA isolation, quality control, library preparation, and RNA sequencing

Subtype-specific GCs or somata were collected in RLT buffer supplemented with 10% b-Mercaptoethanol, then lysed by vortexing for 30-60 sec. RNA was extracted using an Allprep kit (Qiagen), following the manufacturer ’s protocol. Elution was performed in 13 μl RNase-free water, and samples were stored at -80°C. Quality and concentration of extracted RNA was assessed using a Tapestation 2100 with the High Sensitivity RNA kit (Agilent) for samples extracted form sorted somata and using a 2100 Bioanalyzer with the RNA 6000 Pico kit (Agilent) for samples extracted from sorted GCs. Only samples that met quality control (QC) standards were submitted for library preparation and RNA sequencing (RNA-seq). Libraries were prepared from <=2.5ng input RNA using ¼ volume SmartSeqv4 technology (Takara Bio) and sequenced on NextSeq High Output flow cells. Each library resulted in >=40M fragments sequenced (see Supplementary Figure 3a).

### RNAseq data processing and differential expression analysis

The full RN-seq data processing pipeline is available at https://github.com/priyaveeraraghavan/amalgam, schematized in Supplementary Figure 3b. Briefly, adapters were trimmed and low-quality reads were filtered using the Trim-Galore! wrapper for Cutadapt and FastQC. Sequences aligning to rRNA, rodent repeat elements, 7SL or SRP (the RNA component of the signal recognition particle), or the mitochondrial genome were removed using bbsplit.sh from the BBTools suite (Supplementary Figure 3c). Reads were aligned to the GRCm38 genome and to the corresponding Ensembl transcriptome version 101 using STAR (version 2.7.9a) with default parameters.

Gene level differential expression: Transcript abundances were estimated using Salmon (version 1.7.0) alignment mode with [–posBias –numBootstraps 100] flags set. Gene-level differential expression analysis was performed using DESeq2 (version 1.26.0). GC genes were defined as those that were significantly more abundant in GC samples compared to input/GCF samples using a one-sided Wald test with an FDR cutoff of <0.05. When comparing between two GC sample groups, we compare the union of genes that are enriched above GCF. For both GC and soma comparisons, genes were considered significant using an FDR < 0.05 and fold change of >2 for somata and >0 for GCs, unless otherwise stated.

Isoform level differential expression: For 3 ’UTR isoform analysis, only the last 250bp of a transcript are counted and compared between samples, due to 3 ’ sequence coverage bias in the data. A custom gtf was prepared containing these last 250bp (available on github), collapsing transcripts that have overlapping regions. A “UTR group” is defined by the longest UTR represented by the group that has the same/overlapping 3 ’ end. Reads falling within these regions are counted using the featureCounts tool from the subread package. GC 3 ’UTRs are defined as having a log2 fold change >0.5 of GC/GCF with an FDR cutoff at 0.05, more stringently than in the gene level analysis.

Output of differential expression analysis on gene and transcript level is provided in Supplementary Table 2.

### GO term mapping across subcellular compartments and subtypes

GO term enrichment was computed for each GO term and gene set using Fisher ’s Exact Test, using ClusterProfiler (version 4.2.0). For details, see https://github.com/priyaveeraraghavan/amalgam. For GO term dotplots, the compareCluster function was used. For genes differentially abundant between CPN and CThPN GCs, all significantly overrepresented GO terms (FDR < 0.05) in either group were retained. GO terms were simplified with the Wang similarity measure, using a similarity threshold of 0.7. Then, for each group, the - log10(p-value) for each simplified GO term was calculated and displayed in the dotplot. The full list of all enriched GO terms across compartments and sub-types can be found in Supplementary Tables 1 and 4. GO-term chordplot (Fig 2f): For select GO terms identified as enriched in CPN or CThPN GCs, the differentially enriched genes associated with that GO term are connected to the GO term.

### Representation of genes associated with neurodevelopmental diseases in GC-localized transcriptomes

Enrichment of disease-associated genes in each subtype/subcompartment was calculated using Fisher ’s Exact Test, and the resultant odds ratios and 95% confidence intervals plotted. A tabular version of the results can be found in Supplementary Table 3. The following gene sets were used as references:-Disease-associated genes: (1) Schizophrenia-associated rare coding variants identified via whole exome sequencing (WES) with FDR < 0.05 from Singh, et al. 2022^74^; (2) Prioritized/fine-mapped schizophrenia-associated genes defined in Extended Data Table 1 of Trubetskoy, et al. 2022^75^; (3) ASD-associated genes associated with variants that reached exome-wide significance in Satterstrom, et al. 2021^72^; (4) Bipolar-associated protein truncating variants with a Fisher p-value < 0.05 from Palmer, et al. 2022^133^ (NB: for this BipEx dataset, no variant reached exome-wide significance).

-Subcellular/subtype sets: For each of CPN and CThPN subtypes, GC genes were defined as being present in GCs above experimental background. Soma genes were defined as the remaining set of expressed but non-GC genes in the respective subtype.

### Subcellular mapping of WIRS-containing transmembrane and membrane associated proteins

WIRS-containing transmembrane and membrane associated proteins are the set defined by Chen et al.^134^ Supplementary Figure 2.

### 3 ‘UTR motif analysis

De Novo: GC-enriched 3 ’UTR sequences were compared to all expressed 3 ’UTR sequences using STREME (5.4.1) in the -de mode.

Scanning/enrichment of known motifs: The Transite tool^78^ was used to scan for a curated set of 174 RBP-associated, matrix-defined motifs across 3 ’UTRs, and to subsequently calculate enrichment or depletion of motifs in a subset of 3 ’UTRs (e.g. GC-enriched UTRs vs soma). Individual sequence k-mers were counted per UTR using the str_locate_all function from stringr (1.5.0).

## Transcardial perfusion, tissue processing, and immunocytochemistry

Postnatal mice were deeply anesthetized using hypothermia, and transcardially perfused with 3-5 ml of pre-cooled PBS followed by 5 ml pre-cooled 4% paraformaldehyde. Brains were carefully extracted from the skull and postfixed in 4% PFA overnight. The following day, brains were washed in PBS and cryoprotected with 30% sucrose in PBS. Tissue was embedded in O.C.T (#25608-930, Sakura Finetek USA Inc), sectioned using a cryostat (Leica, 50 μm thick coronal sections for ICC), and collected in PBS supplemented with 0.025% Azide. For immunocytochemical labeling, tissue was blocked in PBS supplemented with 0.3% TritonX100, 0.3% BSA, and 2% donkey serum (blockperm) for 1 hour at room temperature, then incubated with primary antibodies (RFP #600-401-379 Rockland; GFP #A-11122 Invitrogen; SatB2 #ab51502 abcam; Tbr1 #ab31940) in blocking buffer on a rocker over night at 4°C. The following day, sections were washed three times in PBS on an orbital shaker, incubated for 2-3 hours at room temperature with isotype-specific, fluorescently-conjugated secondary antibodies (Life Technologies) and DAPI (1:10,000, #D1306 Thermo), washed again three times with PBS, mounted on glass slides (Supefrost, #48311-702 VWR), coverslipped with fluoromount (#0100-01 VWR/Southern Biotech), and sealed with nail polish.

Fluorescence images were acquired using an epifluorescence microscope equipped with a motorized stage and a 10x objective (Nikon NiE). Whole brain sections were imaged using EDF z-stack projections and mosaic image stitching through NIS Elements software (Nikon).

### Quantification of CPN IUE targeting

Images were processed using Fiji software (ImageJ 1.53t). For each mouse, the coronal section closest to the fused anterior commissure was identified. Fluorescently labeled cell bodies in the ipsilateral cortical hemisphere were identified manually, and coordinates were extracted. To account for curvature of the brain surface, respective coordinates were projected onto the dorsal surface and plotted relative to the media-lateral extent of the cortex using Matlab (R2022b Update 5, Mathworks).

### Quantification of CPN midline crossing and contralateral grey matter innervation

Epifluorescence images were processed using Fiji software (ImageJ 1.53t). For each mouse, the coronal section closest to the fused anterior commissure was identified. For quantification of CPN elongation in the corpus callosum and lateral white matter, innervation into the cortical grey matter, and fiber abundance in cortical layer V and II/III, respective regions of interest (ROI, line of 200 pts width) were manually traced as shown by outlines in the schematic insets in Supplementary Figure 11b-e. Briefly, ROIs for “elongation in CC and lateral white matter” started at the midline of the corpus callosum and followed the lateral white matter to its lateral extent. ROIs for “innervation into cortical grey matter” were positioned just dorsal to the lateral white matter. ROIs for “fiber abundance in layers V and II/III” were positioned on the ventral side of layer V and II/III, respectively, as identified in the DAPI channel. The GFP channel of the ROIs was straightened using Fiji ’s standard straightening tool, binarized by applying respective thresholds, binned, and analyzed for mean grey value. Data from individual mice was averaged across groups and reported as mean±sem using R studio (RStudio 2023.03.0, Posit Software, PBC).

### Primary neuronal cultures

CPN were fluorescently labeled and/or targeted for shRNA-mediated knock-down using ipsilateral *in utero* electroporation of respective DNA constructs outlined above at E14.5. At birth (*P*0), cortical projection neurons were collected using the soma preparation protocol outlined above. For experiments in the context of developmental stages, cortical projection neurons were isolated at P1 or P3 to reflect respective gene expression states, using the same protocol. Instead of sorting fluorescence-positive neuronal somata using FACS, the final cell suspension was cultured on pre-coated glass coverslips (50 μg/ml PDL #P0899 Sigma, in ddH2O over night at 37°C, followed by five thorough washes in ddH2O and incubation with 10 ug/ml mouse laminin (#23017015, Thermo) in magnesium- and calcium-free DPBS (#14190-144, Gibco) for 1-2 hours at 37°C) in 24-well plates at standard conditions (37°C, 5% CO2). For immunocytochemistry, cultured primary neurons were washed with PBS 24 hours post isolation and fixed with 4% PFA for 10 min, followed by final washes and storage in PBS supplemented with 0.025% azide. For RNAscope, cultured primary neurons were washed with PBS 24 hours post isolation and fixed with 4% PFA 30 min, followed by two washes in PBS, dehydration in 50%, 70%, and 100% ethanol, and stored in 100% ethanol at -20°C.

### Immunocytochemistry

Coverslips with fixed primary neurons were transferred to 4-well slides and recovered in PBS for 10 min. Cells were permeabilized with 0.2% Triton-X100 in PBS for 5 minutes and washed 3 times over 6 minutes with PBS-T (PBS + 0.02% Tween-20; one quick wash followed by 2x3 min). Samples were blocked in PBS-T supplemented with 3% BSA and 2% donkey serum (blocking buffer) for 30 min before incubation with primary antibodies (GFP #A-11122 Invitrogen; RFP #600-401-379 Rockland; Actin #A5441 Sigma; WAVE1 #ab272916 abcam; Cpeb4 #ab224162 abcam; Rbms1 #ab150353 abcam) in PBS-T+3% BSA overnight at 4°C. The next day, coverslips were washed three times in PBS-T (1x fast wash, 2x 5min), incubated with isotype-specific secondary antibodies (Life Technologies) and DAPI (1:10,000, #D1306 ThermoFisher) for 2-3 hours at room temperature in PBS-T+3% BSA on an orbital shaker, followed by three final washes in PBS-T as described above, mounting on a glass slide using fluoromount (#0100-01, VWR/Southern Biotech), and sealing with nail polish. Fluorescence images were acquired using an epifluorescence microscope equipped with a motorized stage and a 40x oil objective (Nikon NiE, EDF z-stack projections through NIS Elements software (Nikon)), as well as a confocal microscope equipped with a 40x oil objective (LSM880, Zeiss, maximum intensity projections of 10 μm z stack).

### Quantification of protein abundance along longest neurite

Confocal images were processed using Fiji software (ImageJ 1.53t). Only neurons that had a clearly visible and uncrossed primary neurite were included for analysis. Each neuron was manually traced across the maximum extent of the cell body and along the longest neurite, and respective line profiles were plotted using the standard Fiji “plot profile” tool for the GFP, DAPI, Actin, and protein of interest channels. Respective data were averaged across groups and reported as mean ± sem using R studio (RStudio 2023.03.0, Posit Software, PBC).

### Sholl analysis of branching complexity of cultured neurons

Confocal images were processed using Fiji software (ImageJ 1.53t). Only neurons that had a clearly visible and uncrossed primary neurite were included for analysis. The GFP channel of each image was binarized using respective thresholds, and a line ROI was drawn from the center of the cell body to the distal tip of the longest neurite. Sholl analysis (10 μm step size) of the longest neurite was performed using the respective plug-in from Fiji ’s SNT framework^135^. Data were averaged across groups and reported as mean #(intersections) ± sem using Excel.

### RNAscope in cultured neurons

Coverslips with fixed, and dehydrated primary neurons were stored at -20°C. Coverslips were transferred to 4-well slides and rehydrated in a stepwise fashion using 100%, 70%, 50% ethanol for 1 min each, followed by two washes with PBS for 1 min each. For single candidate RNAscope experiments (WAVE1), the ACD Bio chromogenic RNAscope RED kit was used (ACDbio, #322350), following the manufacturer ’s instructions. For multiple candidate RNAscope experiments (WAVE2/3 or Pcdh17/Robo1), the ACD Bio multiplex V2 RNAscope kit was used (ACDbio, #323100), following the manufacturer ’s instructions.

After single candidate or multiple candidate RNAscope detection, coverslips were subjected to subsequent immunocytochemistry to enhance the GFP signal. Briefly, following the final RNAscope wash, coverslips were blocked and permeabilized in PBS supplemented with 0.03% TritonX100, 0.3% BSA, and 2% donkey serum (block/perm) for 30 min, and subsequently incubated with primary antibody (GFP, #A-11122 Invitrogen) in block/perm buffer overnight at 4°C. The next day, coverslips were washed three times for 10 min each in PBS, incubated with isotype-specific secondary antibodies (Life Technologies) and DAPI (1:10,000, #D1306 ThermoFisher) for 2-3 hours at room temperature on an orbital shaker, followed by three final washes in PBS for 10 min each, mounting on a glass slide using Fluoromount (#0100-01, VWR/Southern Biotech), and sealing with nail polish. Fluorescence images were acquired using a confocal microscope equipped with a 63x oil objective (LSM880, Zeiss, maximum intensity projections of 10 μm z-stack).

**Supplementary Table 1:** List of GO terms differentially enriched in either soma or GC compartments for union of CPN and CThPN

**Supplementary Table 2:** Results of differential expression analysis at gene and transcript level, comparing expression across subtypes of cortical projection neurons, compartments, and across developmental stages for CPN.

**Supplementary Table 3:** Cross-referencing subcellular differential gene expression data of this report for CPN and CThPN with previously published, data sets of genes associated with neurodevelopmental and neuropsychiatric disorders.

**Supplementary Table 4:** List of GO terms differentially enriched in GC compartments of distinct subtypes of cortical projection neurons.

**Figure S1:**
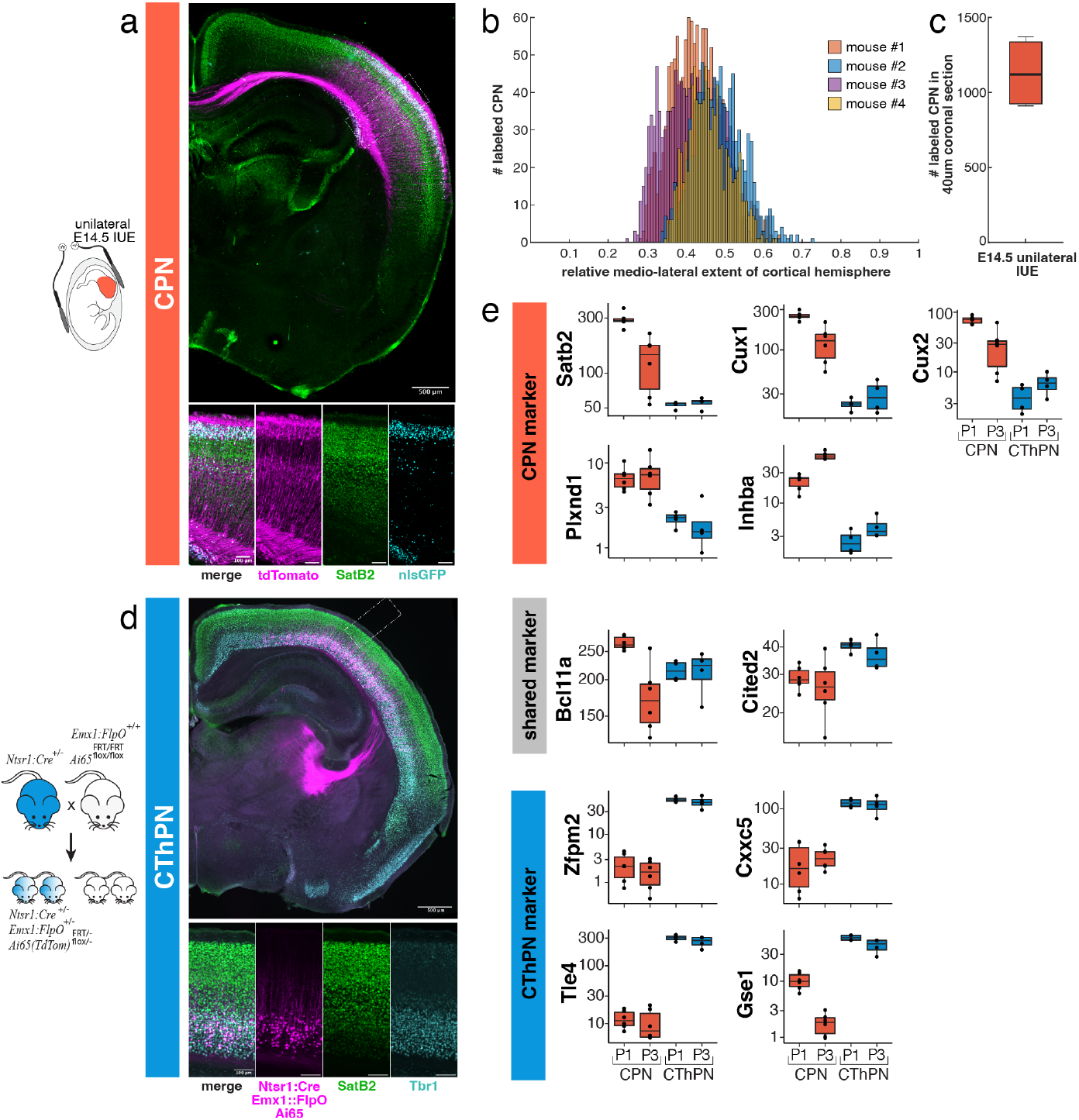
Subtype-specific labeling of CPN and CThPN. **(a)** *In utero* electroporation at embryonic day E14.5 in mice specifically labels CPN of cortical layer II/III. GFP-positive nuclei of CPN are localized mainly to cortical layer II, with limited labeling present in layer III, and are colocalized with the superficial layer marker SatB2 (inset). The multi-cistronically expressed, membrane-targeted tdTomato reveals the entire extent of CPN dendritic processes and axonal projections spanning all cortical layers, as well as across the corpus callosum. **(b)** Unilateral *in utero* electroporation at embryonic day E14.5 results in highly reproducible targeting of CPN in cortical layer II/III in the lateral somatosensory cortex. **(c)** Total number of labeled CPN in a single 40 μm coronal brain section is comparable across mice. **(d)** Intersectional mouse genetics (Ntsr1::Cre+/-, Emx1::FlpO+/-, Ai65(TdTom)FRT/wt, flox/wt) specifically labels CThPN of cortical layer VI. Labeled CThPN projections transverse the internal capsule and innervate the thalamus. TdTom-positive CThPN are colocalized with CThPN marker TBR1 but are negative for superficial layer marker SatB2 (inset). **(e)** RNA expression levels of subtype marker genes from flow cytometry sorted CPN and CThPN somata at developmental ages P1 and P3. CPN have markedly higher expression levels in superficial layer markers, whereas CThPN have appropriate enrichment for layer VI marker genes, as expected. Genes common to both subtypes are expressed at similar levels.

**Figure S2:**
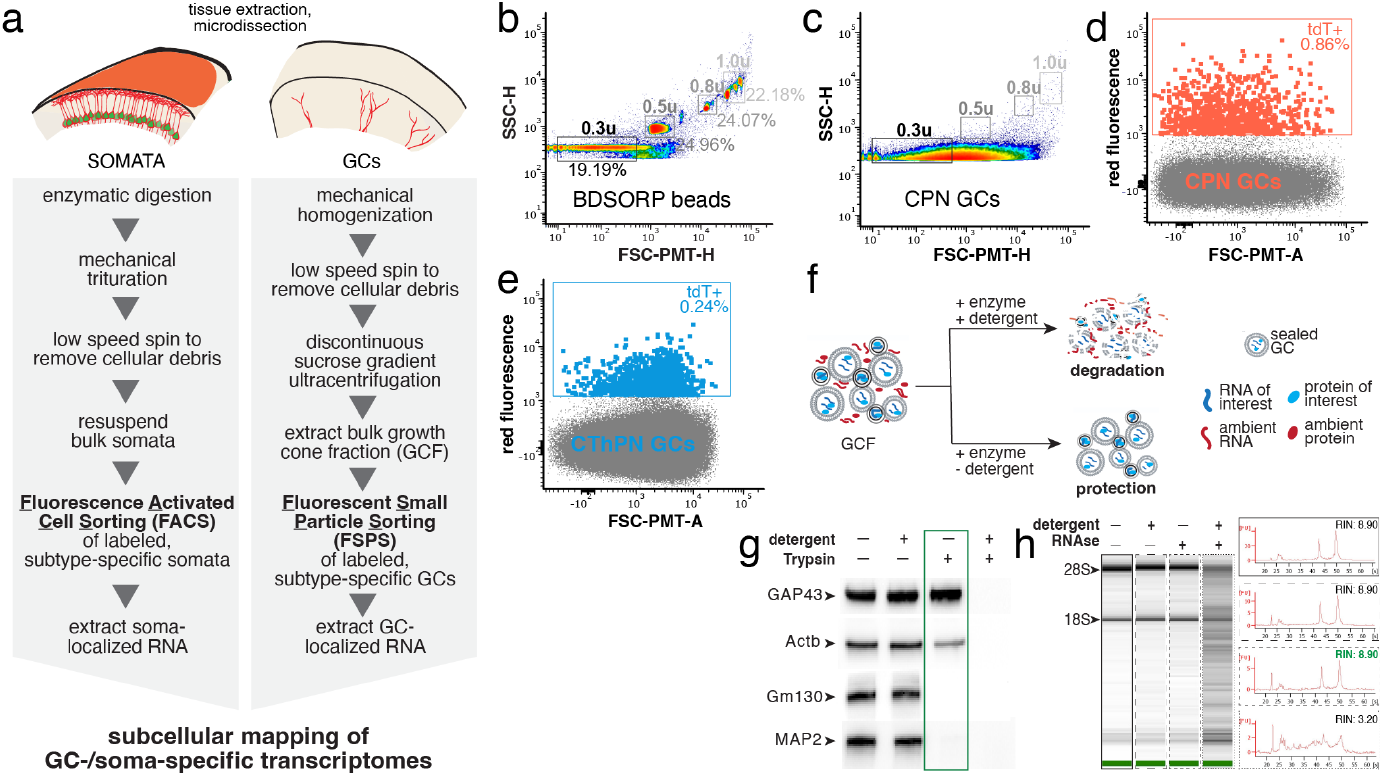
Parallel isolation of subtype-specific somata and GCs from the mouse neocortex. **(a)** Workflow enabling parallel extraction and subtype-specific sorting of CPN- or CThPN-specific somata and GCs. RNA isolated from respective samples sorter-purified by flow cytometry is then used for subcellular mapping of subtype-specific transcriptomes. A detailed description of this protocol can be found in^50^. **(b)** Side scatter over forward scatter data for size-calibrated BD SORP beads run on a flow cytometer in fluorescent small particle sorting (FSPS) configuration. FSPS enables detection, analysis, and efficient sorting of particles in the 100nm-1 μm diameter range. **(c)** Analysis of CPN GCs overlaid on top of BD SORP beads depicted in detail in panel (b). CPN GCs have diameters of ∼200-800 nm, predominantly ∼300-500 nm. **(d/e)** FSPS enables purification of subtype-specifically labeled GCs of CPN (d) and CThPN (f) from respective GCF samples. **(f)** Hydrolysis protection assays are used to assess the presence of intact GC particles with intact membranes, thereby protecting their specific molecular contents from RNase or proteinase digestion in the absence of detergents. **(g)** Western blot of representative proteinase protection assay of CPN GCF detected for multiple subcellular markers. Likely ambient contaminations like GM130 or Map2 are degraded in the absence of detergents (-/+), while GC markers such as GAP43 or Actin are preserved in the absence of detergents (-/+), and are only degraded once GC membranes are permeabilized by detergent (+/+). **(h)** Tape station results of representative RNase protection assay of CPN GC. When the sample is treated with RNase, RNA quality is preserved in the absence of detergents (-/+), and only upon permeabilization of membranes does pronounced RNA degradation occur (+/+).

**Figure S3:**
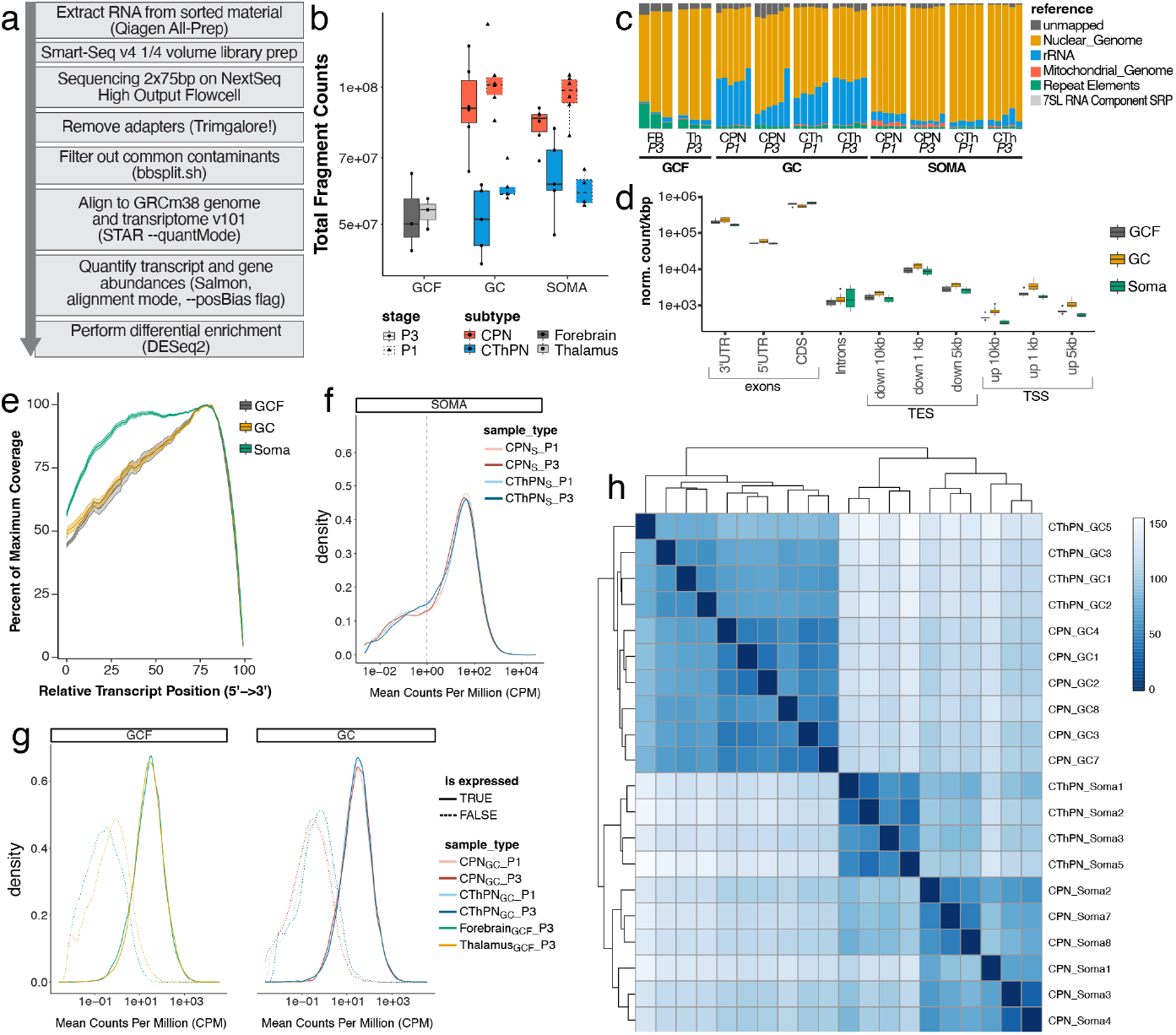
Subcellular RNA sequencing passes quality metrics and samples are highly correlated within compartments and subtypes. **(a)** Schematic of bioinformatics workflow to process transcriptomic data from reads to differential expression between samples. **(b)** Sequencing depth is >40M fragments for all libraries across subtypes and stages, as well as for background samples (GC fraction pre-sort, GCF). **(c)** Alignment of reads to contaminating features and the nuclear genome indicates that most reads, as expected, originate from the nuclear genome and that the sorted GC samples have a higher proportion of reads aligning to rRNAs. **(d)** Distribution of reads to genomic features shows that most reads align to UTRs and coding sequences (CDSs), and not genomic regions downstream of transcription end site (TES) or upstream of transcription start site (TSS), as expected; GC and GCF samples have higher representation in 3 ’UTRs. **(e)** The coverage over 3 ’ ends is higher in GC and GCF samples relative to somata. **(f)** A cutoff of 2 counts per million (CPM) was used for soma samples, since this is the inflection point between signal and noise CPM distributions. **(g)** A CPM cutoff for sorted GC and GCF samples was defined as the 95^th^ quantile of the distribution of CPMs of non-expressed genes (dotted lines), which were determined by the soma cutoff in panel (f). **(h)** Heatmap showing correlation between normalized transcript counts of all pairs of samples.

**Figure S4:**
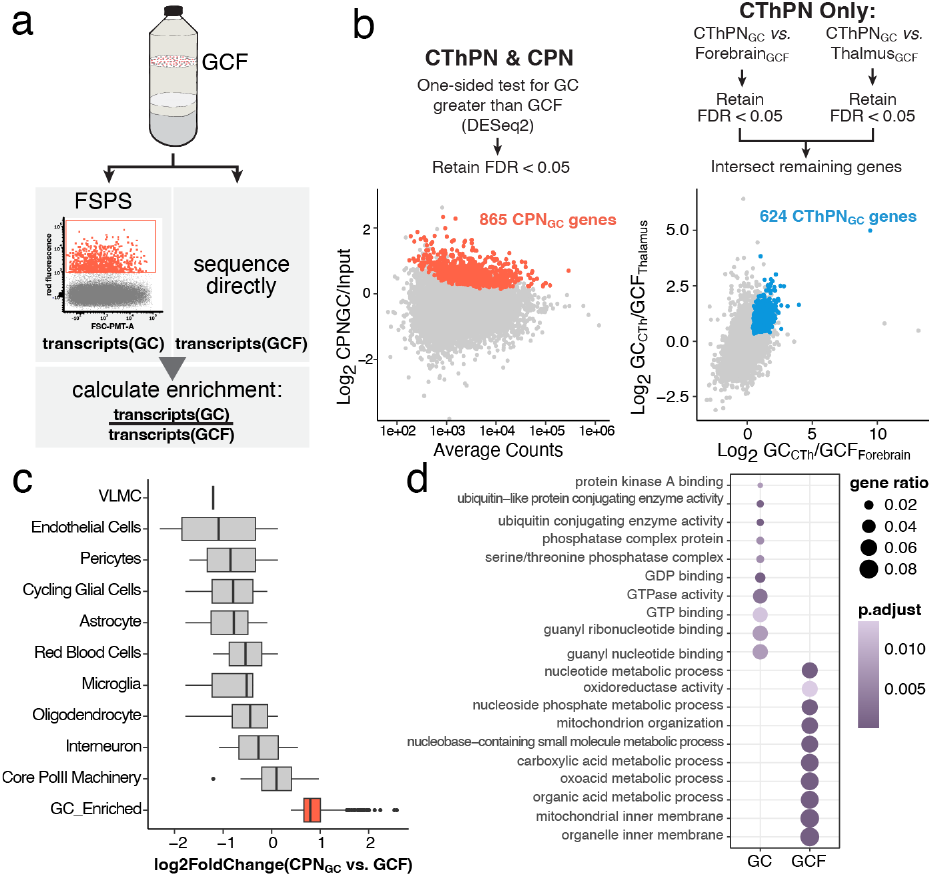
Transcriptome obtained from GC samples is corrected to minimize effect of potential ambient contamination. **(a)** To minimize effects of potential ambient contamination in the GC preparation, transcriptomes obtained from sorted GC samples are corrected for their respective input material, i.e. RNA extracted from the unsorted GCF. **(b)** GC transcripts for CPN and CThPN are defined as those having a higher abundance in sorted GCs than in GCF. **(c)** Correction for input material results in exclusion of likely ambient contaminants, e.g. marker genes for various non-neuronal cell types (grey), while preserving a class of robustly GC transcripts (red). **(d)** GO term enrichment comparing GC genes to likely contaminants from the GCF.

**Figure S5:**
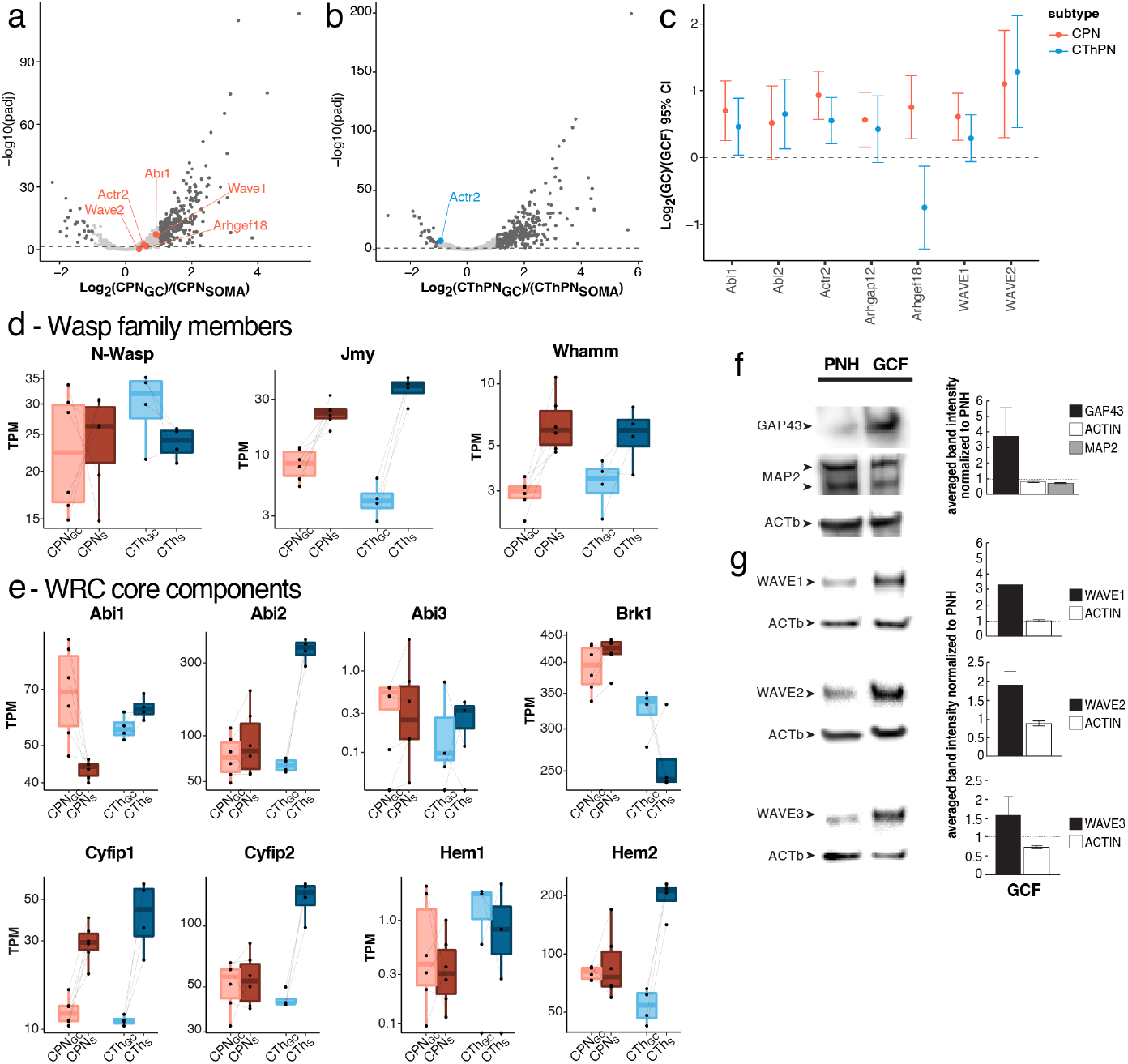
Expression and subcellular transcript localization of WRC core components in CPN and CThPN. **(a)** Volcano plot indicating CPN-specific subcellular transcript localization of core components of the WRC as orange dots (FDR < 0.05). **(b)** Volcano plot indicating CThPN-specific subcellular transcript localization of core components of the WRC as blue dots (FDR < 0.05). **(c)** 95% confidence intervals of log2 fold change GC/GCF for each of the WRC-associated transcripts. **(d)** Transcript abundance (transcripts per million, TPM) of the non-WRC associated members of the Wasp gene family *N-Wasp, Jmy*, and *Whamm* in somata and GCs of CPN (pink/brown) and CThPN (light/dark blue). **(e)** Transcript abundance (TPM) of WRC core components in somata and GCs of CPN (pink/brown) and CThPN (light/dark blue). **(f)** Representative western blot images and quantification of triplicates for each protein ’s abundance of WAVE paralogs WAVE1, WAVE2, WAVE3, and ACTIN in the post nuclear homogenate (PNH) and growth cone fraction (GCF) of CPN. All three paralogs are enriched in the GCF when compared to the PNH.

**Figure S6:**
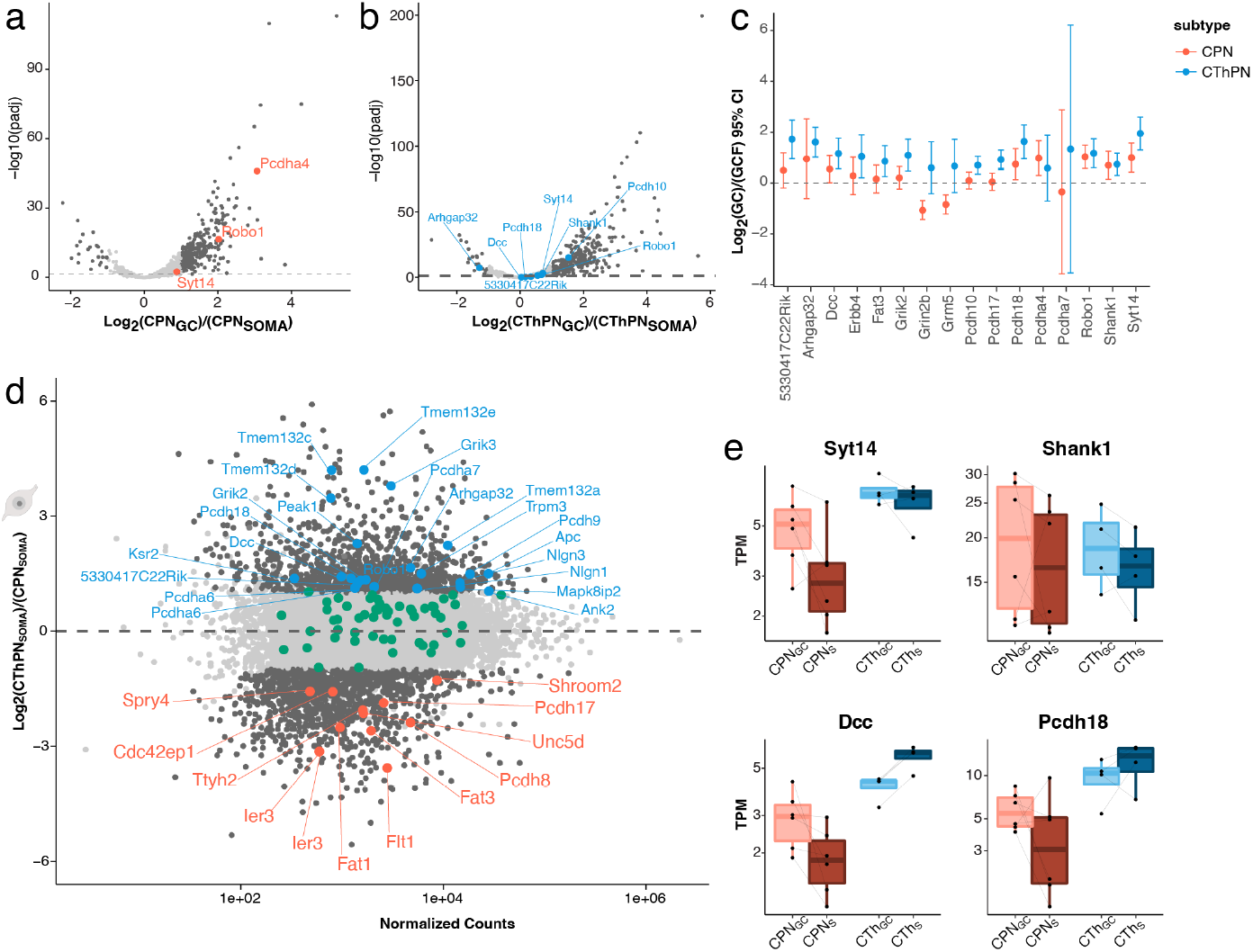
Expression and subcellular transcript localization of WIRS-containing receptors in CPN and CThPN. **(a)** Volcano plot indicating CPN-specific subcellular transcript localization of WIRS-containing receptors as orange dots (FDR < 0.05). **(b)** Volcano plot indicating CThPN-specific subcellular transcript localization of WIRS-containing receptors as blue dots (FDR < 0.05). **(c)** 95% confidence intervals of log2 fold change GC/GCF for each of the WIRS-containing transcripts. **(d)** Differential expression levels (FDR < 0.05, dark grey) of WIRS-containing genes in CPN (orange) vs. CThPN (blue) somata. WIRS-containing genes that are not differentially expressed between subtypes are shown in green. **(e)** Transcript abundance (transcripts per million, TPM) of examples of WIRS-containing receptors with GC-enriched transcript localization in GCs and somata of CPN (pink/brown) and CThPN (light/dark blue).

**Figure S7:**
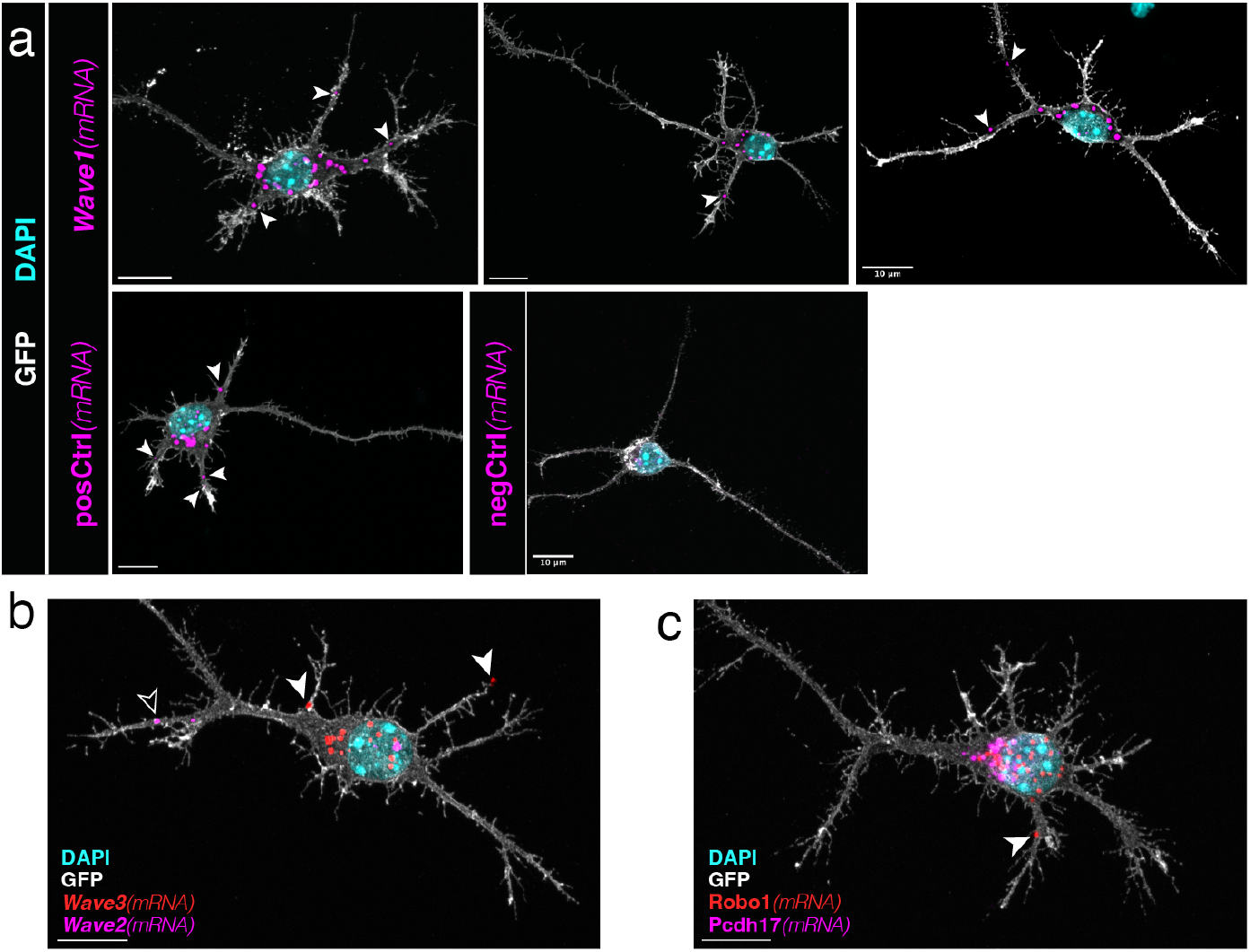
Single molecule FISH of *Wave1* and the WIRS containing receptors *Robo1* and *Pcdh17* in primary cultured CPN. **(a)** Maximum intensity projection (10 μm) of representative RNAscope confocal image of *Wave1*, the respective positive control (posCtrl, targeting *Ppib*), or negative control (negCtrl, targeting *DapB*) in GFP-positive cultured CPN. Solid arrowheads indicate RNAscope puncta localized to proximal or more distal neurites. **(b)** Maximum intensity projection (10 μm) of representative RNAscope confocal image of the *Wave* paralogs *Wave2* (magenta) and *Wave3* (red) in GFP-positive cultured CPN. Solid arrowheads indicate *Wave3* RNAscope puncta, while open arrow heads indicate *Wave2* RNAscope puncta localized to proximal or more distal neurites. **(c)** Maximum intensity projection (10 μm) of representative RNAscope confocal image of the WIRS receptors *Pcdh17* (magenta) and *Robo1* (red) in GFP-positive cultured CPN. Solid arrowhead indicates *Robo1* RNAscope puncta localized to proximal CPN neurite. Similar distribution of puncta were observed for 6-7 neurons per condition.

**Figure S8:**
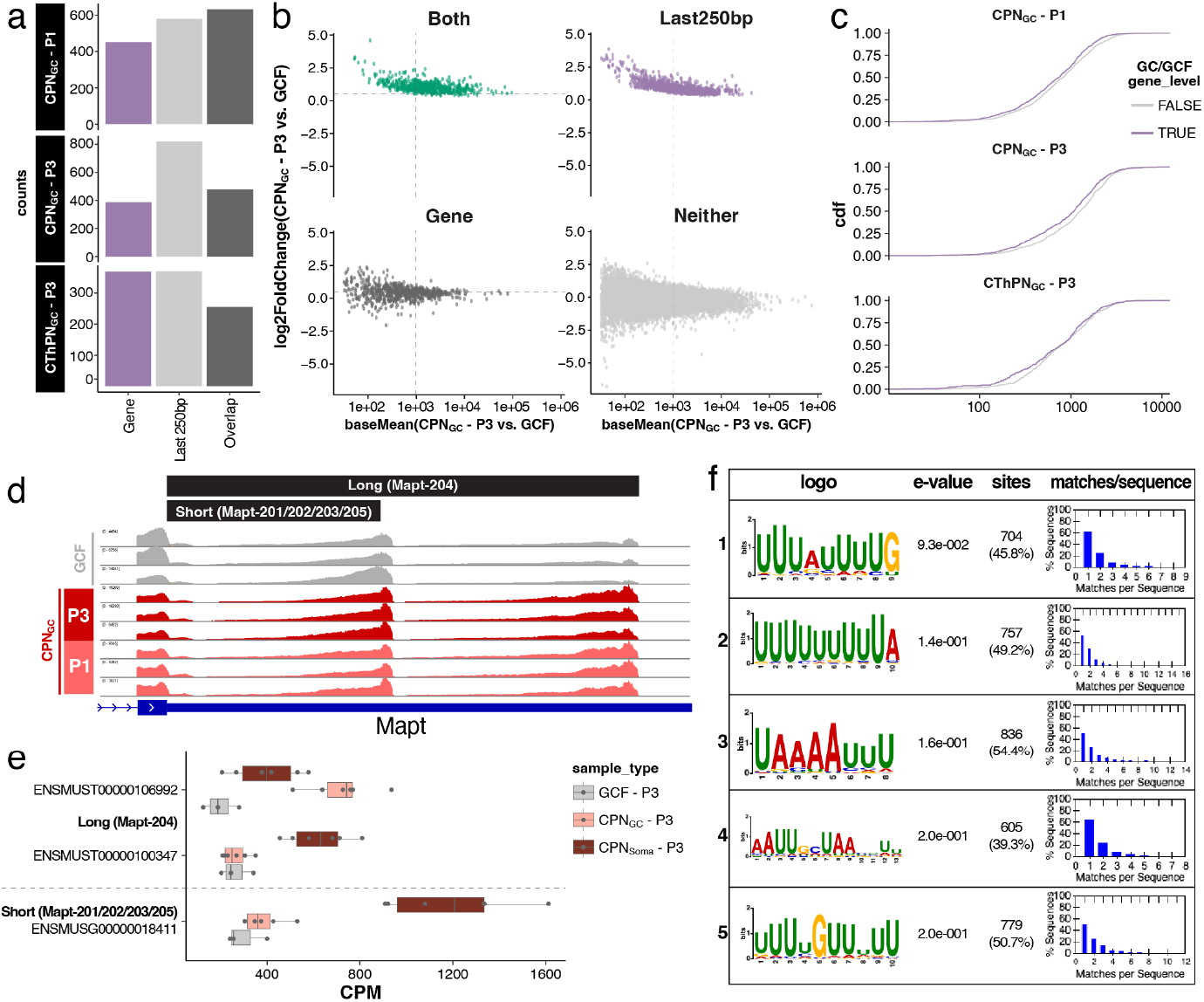
3 ‘UTR isoform analysis identifies enrichment of poly-U motifs in GC transcriptomes over those of somata. **(a)** Number of genes called by the gene-level approach for quantifying GC genes, last-250bp approach for quantifying GC 3 ’UTRs, or both. **(b)** MA plots depicting the log2 fold change and abundance of GC transcripts identified by the last-250bp approach. Each plot represents a subset of all transcripts: genes that are enriched in the last-250bp (purple), genes that were previously identified (black), isoforms of genes that were identified by both approaches (green), and as a control, isoforms of genes that were identified in neither approach (grey). Genes identified by only the gene-level approach (black) are generally of lower abundance. Last-250bp approach (purple) enables identification of GC-specific isoform variants of higher abundance genes. **(c)** GC 3 ’UTRs identified only in the last-250bp approach are on average longer (cdf – cumulative distribution function) (d) As an example, *Mapt* has one long 3 ’UTR enriched in GCs, but a shorter isoform not enriched in GCs, as seen by the read pileups; quantified in **(e). (f)** De novo motif analysis of GC-enriched 3 ’UTRs identifies multiple distinct versions of poly-U rich motifs preferentially detected in the GC compartment.

**Figure S9:**
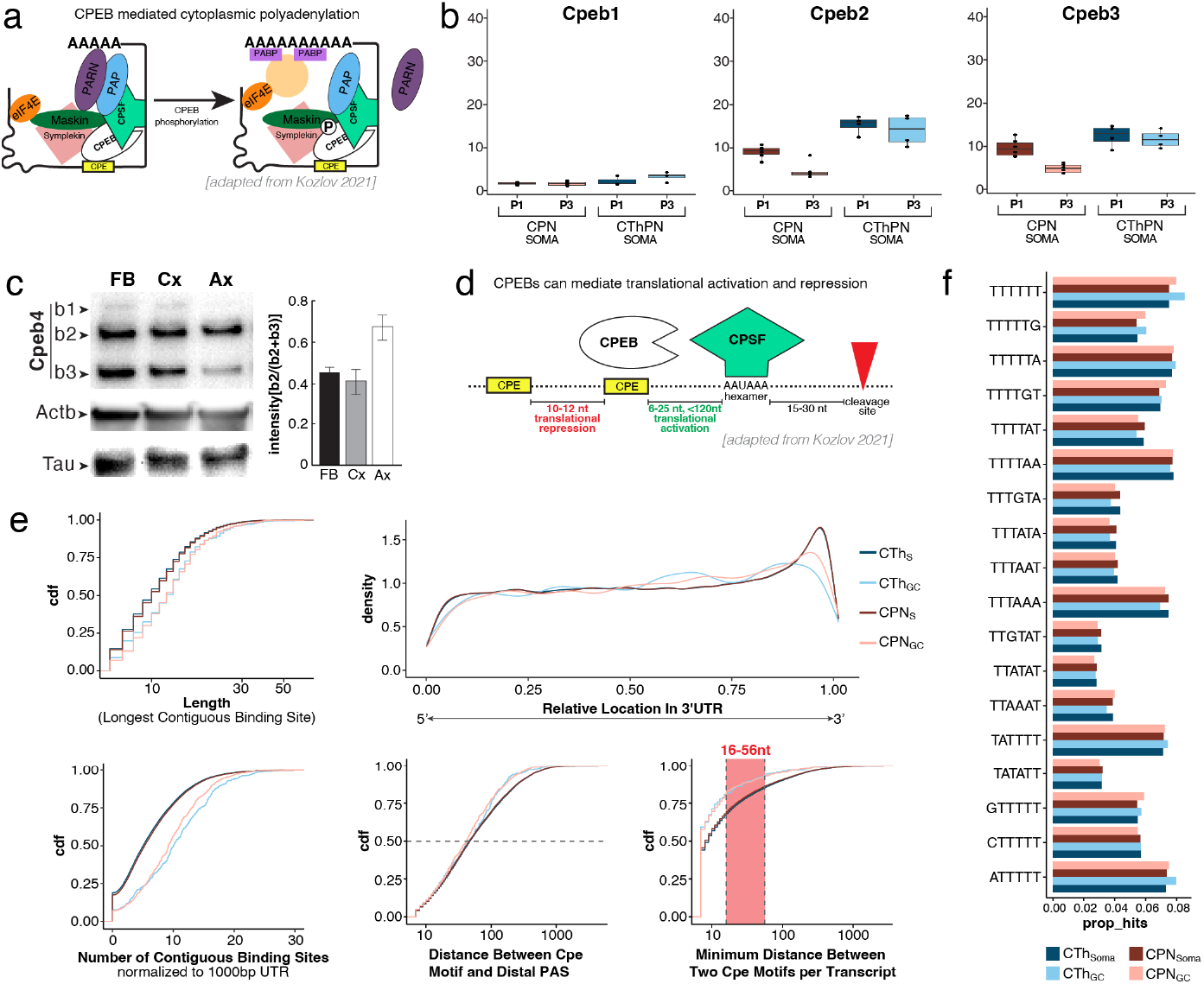
CPE location and spacing in GC 3 ’UTRs is consistent with baseline translational repression of targets. **(a)** CPEBs bind to the cytoplasmic polyadenylation element (CPE) located in the ‘3-UTR of transcripts and complex with factors that control polyA tail length and translation. Cue-induced CPEB phosphorylation mediates polyA polymerase-mediated lengthening of the poly-A tail in the cytosol, enabling eiF4E interaction with PABP, and thereby initiation of translational activation of the respective transcripts [adapted from^96^]. **(b)** Boxplots of expression levels in TPM of the three other *Cpeb* paralogs in CPN (pink/orange) and CThPN (light/dark blue) somata at P1 (brown, dark blue) and P3 (pink, light blue). **(c)** Representative western blot of protein samples obtained from whole forebrain (FB), micro-dissected cortex (Cx), and micro-dissected CPN axon bundles at corpus callosum (Ax), labeled for CPEB4, ACTIN, and TAU. Labeling for CPEB4 reveals 3 distinct bands, which are labeled as b1 (highest MW), b2 (middle MW), and b3 (lowest MW). Quantification of signal intensity in Cpeb4 b1-b3 from triplicates reveals relative enrichment of signal for micro-dissected axons to the middle band (b2). **(d)** Depending on the exact number and relative positioning of CPE motifs within the 3 ’-UTR, CPEBs can mediate either translational activation (as schematized in (a)) or translational repression [adapted from^96^]. **(e)** Quantification of length (top left), number (bottom left), density (top right), distance from the polyadenylation signal (PAS) (bottom middle), and minimum distance (bottom right) between starts of CPE motifs in 3 ’-UTRs enriched in GCs (orange, light blue) and somata (brown, dark blue) of CPN (orange/brown) and CThPN (light/dark blue), cdf – cumulative distribution function. **(f)** Relative proportions of distinct variations of Cpeb4 motif, detected in 3 ’UTRs enriched in GCs (orange, light blue) and somata (brown, dark blue) of CThPN (light/dark blue) and CPN (orange/brown).

**Figure S10:**
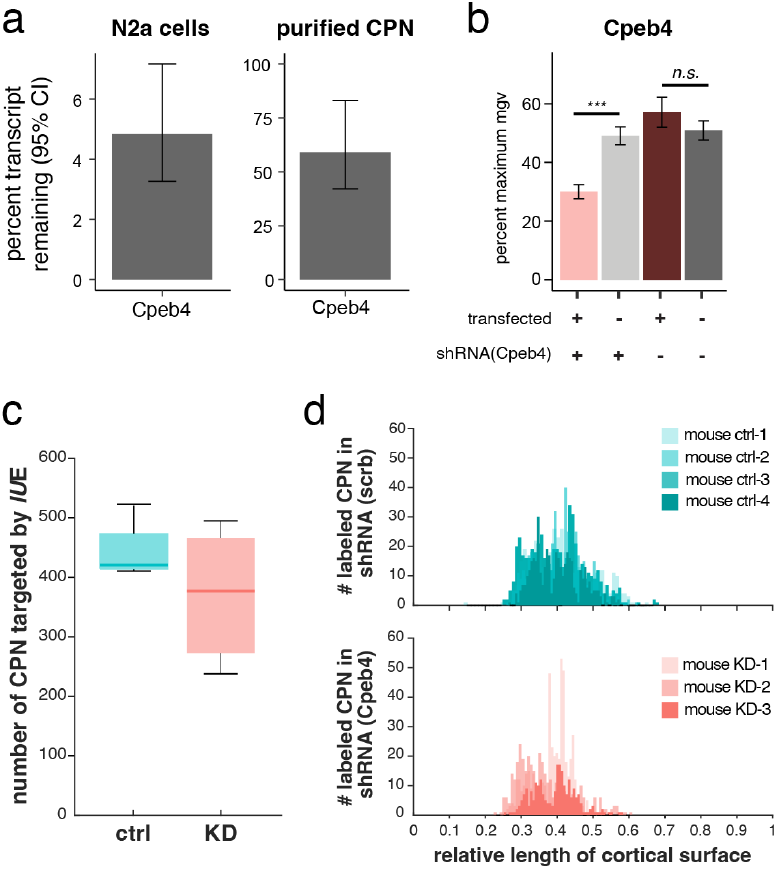
Validation of shRNA-mediated *Cpeb4* knockdown. **(a/b)** Efficiency of shRNA-mediated knockdown of *Cpeb4*. (a) Knockdown efficiency quantified via qPCR in transfected N2a cells and in *in utero* electroporated and FACS-purified CPN. (b) Knockdown efficiency assessed via immunohistochemistry in transfected N2a cells. **(c)** Total number of labeled CPN in a single 40 μm coronal brain section of mice *in utero* electroporated with a scrambled control construct (ctrl) or an shRNA targeting *Cpeb4* for knockdown (KD). **(d)** Unilateral *in utero* electroporation at E14.5 with a scrambled control construct (ctrl, upper panel, n = 4), or shRNA targeting *Cpeb4* (KD, lower panel, n = 3) results in reproducible targeting of CPN in cortical layer II/III in the lateral somatosensory cortex.

**Figure S11:**
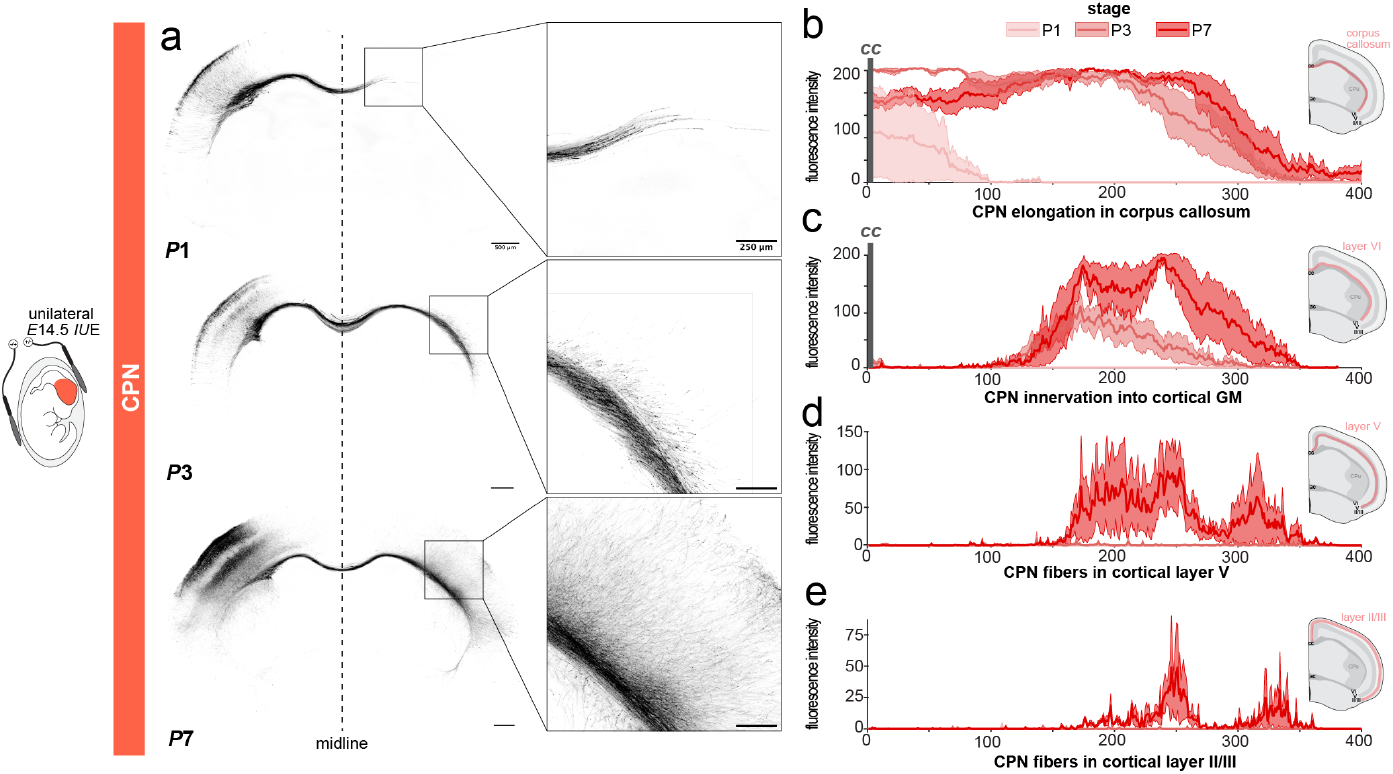
CPN circuit formation occurs during the first postnatal week. **(a)** Somatosensory area CPN extend their axons across the midline around P1, innervate their homotopic target area in the contralateral cortex around P3, and collateralize within the cortical grey matter and likely start formation of synapses around P7. **(b-e)** Quantification of labeled axons/collaterals in progressively entered regions of interest (highlighted in schematic insets to the right, n = 3 for each developmental time), focusing on (b) CPN axon/collateral elongation in the subcortical white matter, (c) innervations into cortical grey matter (GM), (d) axon/collateral density in layer V or (e) in layers II/III at P1 (light pink), P3 (dark pink), and P7 (red).

**Figure S12:**
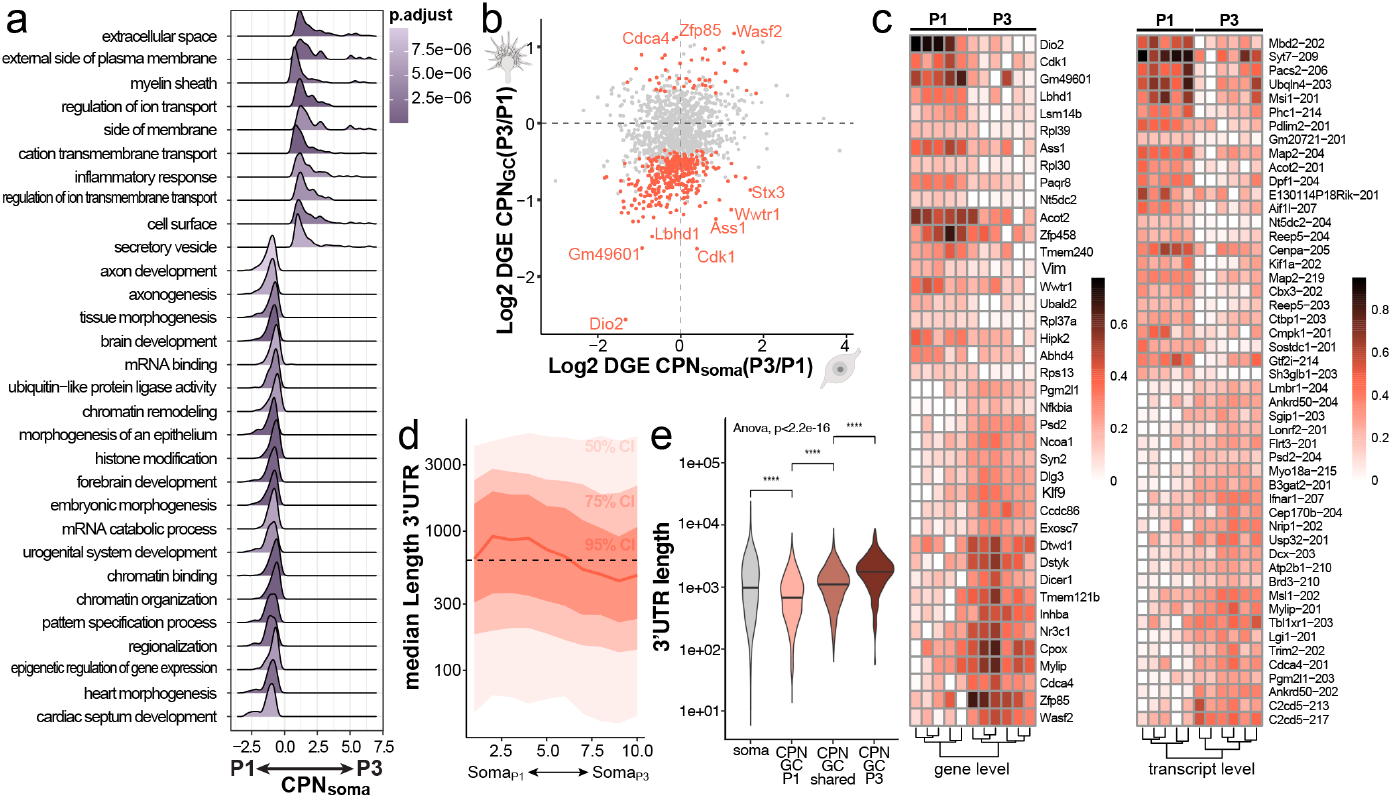
Localized transcriptomes change dynamically from P1 to P3, as CPN switch from axon elongation to grey matter innervation and synapse formation. **(a)** Gene set enrichment analysis for CPN somata comparing P1 and P3. **(b)** Quadrant plot of differential gene expression by CPN at P3 *vs*. P1, and transcript localization by subcellular compartments: in somata (x axis) and GCs (y-axis). **(c)** Scaled transcript abundance of most significantly regulated CPN_GC_ genes comparing P1 and P3, analyzed at gene level (left) and transcript level (right). **(d)** Median length of 3 ’UTR for transcripts detected in CPN and enriched at P1 or P3. Color ribbons indicate 50%, 75%, and 95% confidence intervals (CI). **(e)** Length of 3 ’UTRs of CPN_GC_ genes increases from developmental stage P1 to P3.

**Figure S13:**
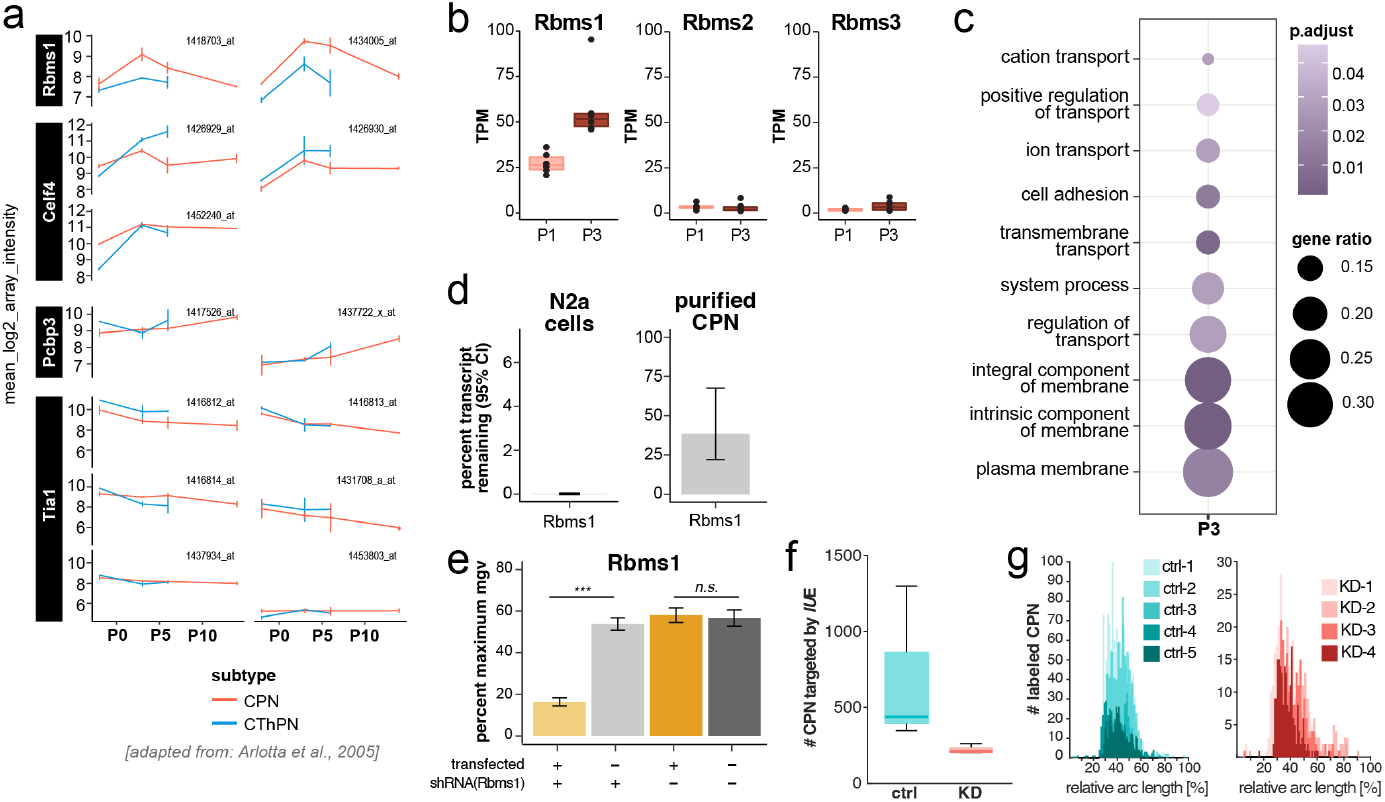
The RBP *Rbms1* increases transcript levels from P0 to P5, and is likely involved in localization and stabilization of CPN_GC_ transcripts. **(a)** Micro array data for RBPs *Rbms1, Celf4, Pcbp3*, and *Tia1* at P0, P5, and P10, analyzed for CPN (red) and CThPN (blue), from Arlotta et al. 2005^51^. **(b)** Boxplots highlighting RNA expression changes for the paralogs *Rbms1, Rbms2*, and *Rbms3* in CPN somata at P1 (pink) and P3 (brown). **(c)** GO term enrichment for subset of CPN_GC_ transcripts containing RBMS1 binding motifs at developmental stage P3. **(d/e)** Efficiency of shRNA-mediated knockdown of *Rbms1*, assessed via (d) qPCR in transfected N2a cells and *in utero* electroporated and FACS-purified CPN, as well as (e) via immunocytochemistry in transfected N2a cells (e). **(f)** Total number of labeled CPN in the cortex of a single 40 μm coronal brain section of mice electroporated with a scrambled control construct (ctrl) or a shRNA targeting *Rbms1* (KD). **(g)** Unilateral *in utero* electroporation at E14.5 with a scrambled control construct (ctrl, left panel, n = 5), or shRNA targeting *Rbms1* (KD, right panel, n = 4) results in comparable positioning of targeted CPN in cortical layer II/III in the lateral somatosensory cortex. The number of labeled CPN was higher in control mice, compared to KD mice.

